# The contribution of recent and historical demographic histories to genomic diversity and conservation status in plant species

**DOI:** 10.64898/2026.06.24.734111

**Authors:** Tongzhou Tao, Pan Li, Yuxiang Zhu, Sining Zhang, Mingming Zhang, Martin Lascoux, Jun Chen

**Author notes:** **Corresponding author**: Jun Chen, **Email:**.

## Abstract

Demographic factors are intrinsically crucial to evaluate species’ extinction risk. However, measuring them remains difficult and time-consuming and the use of genomic summary statistics has been advocated to assess the conservation status of a species. In the present study, we estimated (i) the census number (*N_c_*), (ii) effective population size (*N_e_*) over three different time periods, recent, historical and ancient, (iii) neutral genetic diversity (*π*_4_), and (iv) a measure of the efficacy of purifying selection (*π*_0_⁄*π*_4_) for 101 plant species using population genomic sequencing data. Twenty-one species are from the Plant Species with Extremely Small Populations (PSESP) program of SW China. Threatened species exhibited significantly lower *N_e_*, *N_c_*, *π*_4_, and weaker purifying selection, but had a higher *N_e_*/*N_c_* ratio than non-threatened ones. *N_c_* was the main determinant in identifying conservation status, and contemporary neutral genetic diversity was predominantly influenced by historical *N_e_*. In the absence of demographic information, genetic parameters are a good proxy of conservation status, likely because currently threatened species also had a low historical population size. In summary, our findings suggest that direct estimates of *N_c_* are more useful than *π*_4_, although the latter remains a valuable conservation indicator. Hence, efforts such as the PSESP should be extended.

**Significance Statement:** Extinction is essentially a demographic process. However, evaluating the risk of extinction of species based only on demographic parameters remains a difficult task, and today only 18% of plant species are included in the International Union for Conservation of Nature (IUCN) Red List of Threatened Species. Here, we use demographic data and whole genomic data from 101 plant species to show that while census number remains the best predictor of extinction risk, genomic based parameters provide good alternatives. More specifically, current neutral nucleotide diversity and historical effective population size are good predictors of the extinction risk of plant species.

## Introduction

Since Lande (1988) seminal paper there have been countless discussions on the relative importance of genetics and demography in conservation policies (*1–4*). Lande’s focus was on species extinction and his point was rather straightforward. Since extinction is essentially a demographic process, then demography ought to be the leading principle when establishing conservation policies aiming at avoiding extinction. This is not to say that genetics does not matter, but genetic variation is, in most cases, more important for the long-term well-being of the species than for its short-term survival. In the short-term demographic stochasticity and habitat destruction are more important. This was recently supported by a large-scale survey on the contribution of historical processes to extinction risk in placental mammals (*5*). The authors used genomic data to estimate “historical” effective population size (*N_e_*) and showed that species with smaller “historical” *N_e_* were more likely to be classified as threatened. However, as they pointed out “Ecological (physiological, life-history and behavioral) variables were the best predictor of extinction risk”.

Yet, one of the arguments put forward by conservation geneticists has been that, for most species, estimating population size or characterizing life history traits is generally difficult and that sequencing genomes is today incomparably easier and can provide proxies of population size and an indirect approach to estimate extinction risks (*6*). Since the current IUCN classification of the conservation status of species does not use genetic criteria, this has led to calls “for including genomic information in assessments of the conservation of species” (*7*).

If genomic information is to be considered, which parameters should be retained? One of the most often cited is the “effective population size”, a fundamental concept in population genetics that is directly linked to genetic diversity (*4, 8, 9*). In a Wright-Fisher population, *N_e_* will be equal to the census number but since populations rarely strictly adhere to all requirements of the Wright-Fisher model, the effective population size will rarely be equal to the census number and is generally much smaller (*10*). It is important to be aware that estimates of *N_e_* only provide indirect information on past changes in census number. Also, if a species has experienced a recent and severe decline in population size then its ancestral effective population size can be larger than its current census number as was recently observed in placental mammals (*5*). Various approaches can be used to reconstruct changes in *N_e_* over different time periods (*11*): methods based on linkage disequilibrium retrieve information on past *N_e_* over a few hundred generations, while methods based on the site frequency spectrum (SFS), or on the distribution of heterozygosity along the genome retrieve *N_e_* trajectories across much deeper times but have a low resolution for more recent ones. Which *N_e_* should be used in conservation genetics or, stated differently, which *N_e_* is the most informative on conservation status and extinction risk?

In the present study we compiled population genomic data from 101 plant species with diverse ecological and genetic features, and more importantly, with highly variable distribution ranges and population sizes (**Fig. 1**). For all species, we estimated census number and population genetics parameters, especially three *N_e_* estimates summarizing ancient, historical, and recent changes of population size. With 74 species of known conservation status, we assessed which genetic or ecological features, particularly which *N_e_* estimates, are the most influential, and predicted the extinction risk for 27 species with missing conservation status.

**Figure 1.**
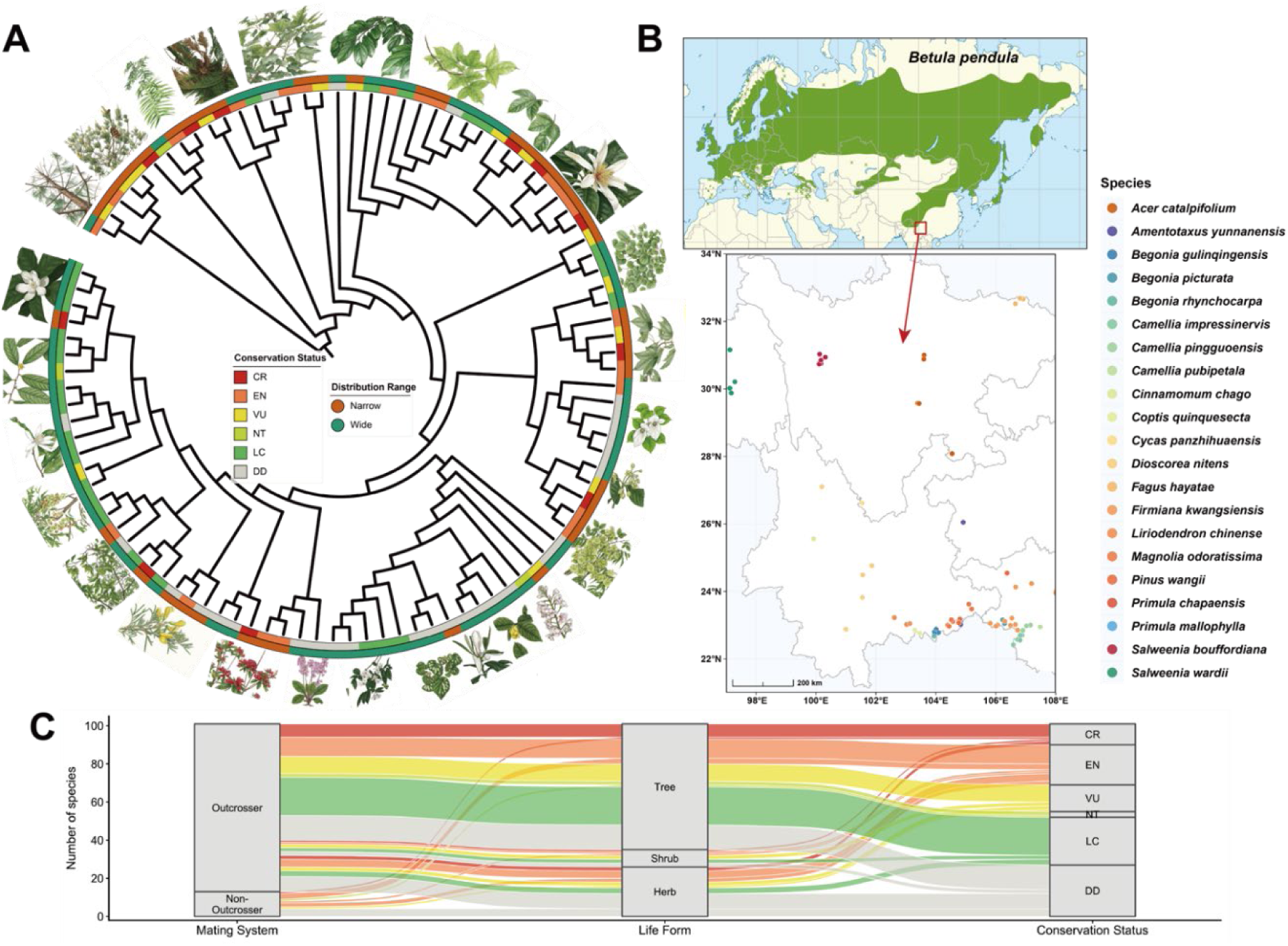
Fig. 1 101 plant species with diverse ecological features. (**A**) The phylogenetic tree relating the 101 species used in the present study. Colors in the inner layer indicate the conservation status. Colors in the outer layer highlight the 41 Chinese species with very limited distribution ranges and small population sizes; (**B**) Examples of distribution ranges of the 21 species with limited distribution ranges in southwestern China and a widely distributed species, *Betula pendula* in Eurasia (source from: https://www.euforgen.org/species/betula-pendula); (**C**) Summary of life form, climate zone, and conservation status for the 101 species.

## Results

Our data comprises 1,933 individual genomes from 101 species across 66 genera and 45 families with varying distribution ranges and ecological features (**Fig. 1**; See details in **Table S1**). Based on their conservation status, all 101 species were grouped into three categories: 1) 49 ‘threatened’ species; 2) 25 ‘non-threatened’ species; and 3) 27 ‘data deficient’ species (DD; See more in **M & M**).

### Threatened species have smaller N_c_ and N_e_

Firstly, a linear correlation model was established between GBIF occurrence records and population census size for tree and non-tree species separately on 44 species for which direct counts of population sizes were available (p-value < 0.001, adjusted R^2^ = 0.85 and 0.51, respectively. **Fig. S1**). This model was then used to estimate population census sizes (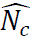) for the remaining 57 species (**Fig. S2**). 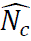 of the threatened group ranged from 34 in *Camellia pingguoensis var. terminalis* (endemic to Daxin County in Guangxi) to 830,000 in *Puya raimondii* (endemic to Bolivia and Peru) with the median equal to 2,000 (**Fig. 2A**). Species of the non-threatened group had a significantly larger median, 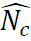, equal to 939,248 (Bonferroni corrected p-value = 1.44×10^-11^), with values ranging from 28,251 in *Gymnocarpos przewalskii* to 2,169,948,935 in *Populus tremula*. Also, the median of group DD’s 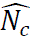 was larger than in the threatened group (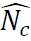 median =107,557, p-value = 7.32×10^-7^).

**Figure 2.**
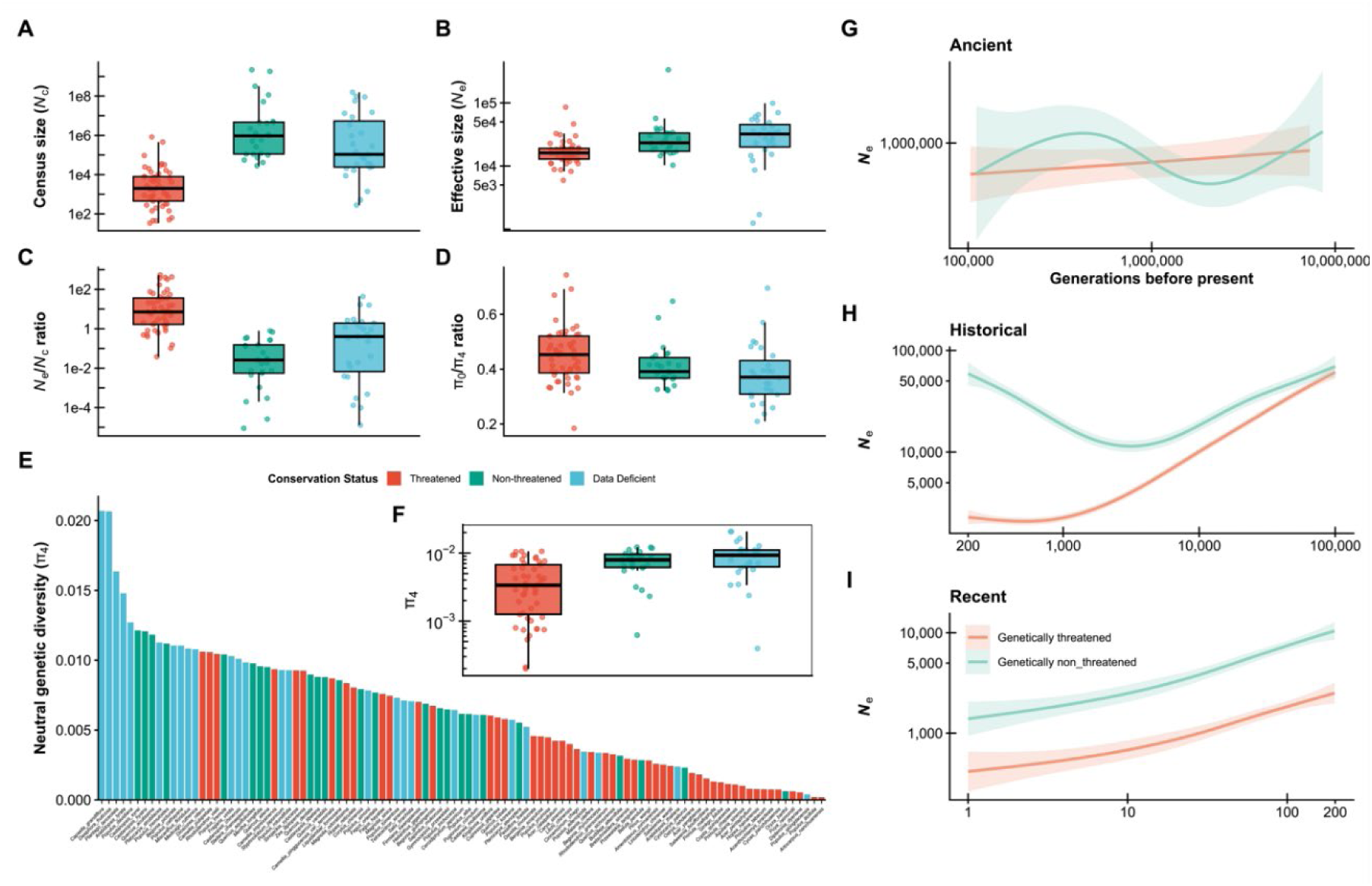
The distributions of population sizes, genetic diversity, and efficacy of selection for three conservation groups. **(A)** Proxy of census size (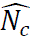); **(B)** Harmonic mean of historical effective population size (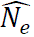) over the generation time interval [200, 10^5^]; **(C**) 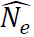/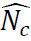 ratio (*N_e_* was derived for historical period estimated by SMCPP); **(D)** *π*_0_⁄*π*_4_ ratio, a measure of the efficacy of selection; **(E)** Pairwise nucleotide diversity at 4-fold sites (*π*_4_) for all 101 species; (**F**) The boxplot of *π*_4_ for three groups; For (**A-F**), threatened species are highlighted in red, non-threatened species are colored in blue, and Data Deficient species are shown in green. (**G-I**) The regression splines of ancient (EPOS), historical (SMCPP) and recent (GONE2) *N_e_* trajectories for 49 threatened species and 3 species (*Populus qiongdaoensis*, *Begonia rhynchocarpa*, *B. picturata*) that were predicted as threatened in ‘randomForest’ analysis in red curves and 25 non-threatened in green curves, respectively.

Next, we estimated effective population size, 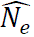, over three different time scales. An ancient 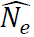 averaged across the time interval [10^5^, 10^7^] was calculated using the SFS-based method, EPOS (*12*) (**Fig. S3A**). For the threatened species, 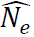 varied from 317 in *Malania oleifera* to 13,617,552 in *Coptis quinquesecta* with a median significantly smaller than for non-threatened species (334,000 *vs* 1,190,000, p-value = 0.01). 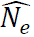 for non-threatened species varied from 1,522 in *Oryza barthii* to 5,189,251 in *Castanopsis eyrei*. The median 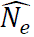 of DD species was equal to 800,000, with the smallest value equal to 31,300 in *Arabidopsis thaliana* and the largest equal to 8,348,156 in *Begonia picturata*. Second, a historical 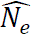 in the generation time interval [200, 10^5^] was obtained with the SMC-based method SMCPP (*13*) (**Fig. 2B**). Like for the previous time period, the threatened group had the smallest 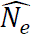 (median = 16,079) compared to the non-threatened (median = 23,274, p-value = 1.78×10^-3^), and group DD (median = 32,065, p-value = 1.25×10^-4^). Third, recent 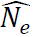, covering the generation time interval [1, 200] was estimated using the LD-based method implemented in GONE2 (*14*) (**Fig. S3B**). In this case, the threatened group had the smallest value, 24 in *Acer yangbiense*, and the largest value, 16,023, in *Dipteronia dyeriana* with a median of 1,652, which again was significantly smaller than in the non-threatened group (varying from 142 to 13,156 with the median equal to 3,611, p-value = 0.03). In this time interval, group DD had a slightly smaller median 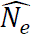 than non-threatened species (from 31 to 116,438 with the median equal to 3,462). In summary, *N_e_*-curves for threatened and non-threatened differed significantly during the historical ([200, 10^5^]) and recent time intervals ([1, 200], **Fig. 2G** **− I and Fig. S4**). In particular, *N_e_* recovered between 200 and 5,000 generations ago after population reduction in non-threatened species but not in threatened species.

Finally, we calculated the ratio *N_e_* /*N_c_* for each species (**Fig. 2C**). *N_e_* /*N_c_* indicates the magnitude of recent demographic change with a large value reflecting an abrupt population decline. Considering historical *N_e_* (in the time interval [200, 10^5^]), 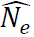/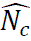 of the threatened group ranged from 535 in *C. pingguoensis var. terminalis* to 0.04 in *Quercus lobata* (endemic to California) and the median equaled 7.17. Species of the non-threatened group had a significantly smaller median 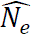/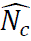, equal to 0.03 (p-value = 5.07×10^-11^), with a minimum value of 8.88×10^-6^ in *Betula pendula* and a maximum value of 0.79 in *Corylus chinensis*. Group DD also had a smaller 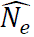/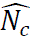 than the threatened group (median = 0.40, p-value = 1.92×10^-5^). The 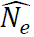/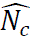 ratios based on ancient and recent *N_e_* estimates were also higher in threatened than in non-threatened species (**Fig. S3**).

### Genetic diversity is determined by both genetic and ecological features

The neutral genetic diversity, *π*_4_, varied 104-fold between the least and the most diverse species (**Fig 2D**). In general, the threatened species had the lowest genetic diversity (median *π*_4_ = 0.0036) but highest accumulated deleterious mutations (median *π*_0_⁄*π*_4_ = 0.45, **Fig. 2D-F**). The median *π*_4_ was more than twice as large in the non-threatened and almost tripled in DD species (0.0079 and 0.0093, respectively; p-value = 8.85×10^-4^). As expected, purifying selection was more efficient in the latter two groups than in threatened species (median *π*_0_⁄*π*_4_ = 0.39 and 0.37, respectively; p-value = 3.57×10^-3^).

With highly correlated features removed (**Fig. S5 & S6.** See all features in **Table S1**), we then evaluated the contributions of the remaining 12 features to species’ neutral genetic diversity using a Bayesian mixed model with the phylogenetic tree considered as a random effect. With a moderate phylogenetic signal (*λ* = 0.408, 95% CI: 0.003-0.909), the historical 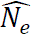 (in the time interval [200, 10^5^]) exerted the largest effect (0.561, 95% CI: 0.426-0.701) on genetic diversity among fixed factors: as expected, species with large historical 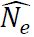 maintained significantly higher genetic diversity. Mating system had the second-largest effect (0.553, 95% CI: 0.197-0.903), with outcrossing species harboring significantly higher genetic diversity than partially or strictly selfing species. GC content, 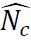, and ancient 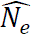 (in the time interval [10^7^, 10^5^]) also had significant and positive effects on genetic diversity (0.325, 0.251 and 0.214, respectively; **Fig. 3**).

**Figure 3.**
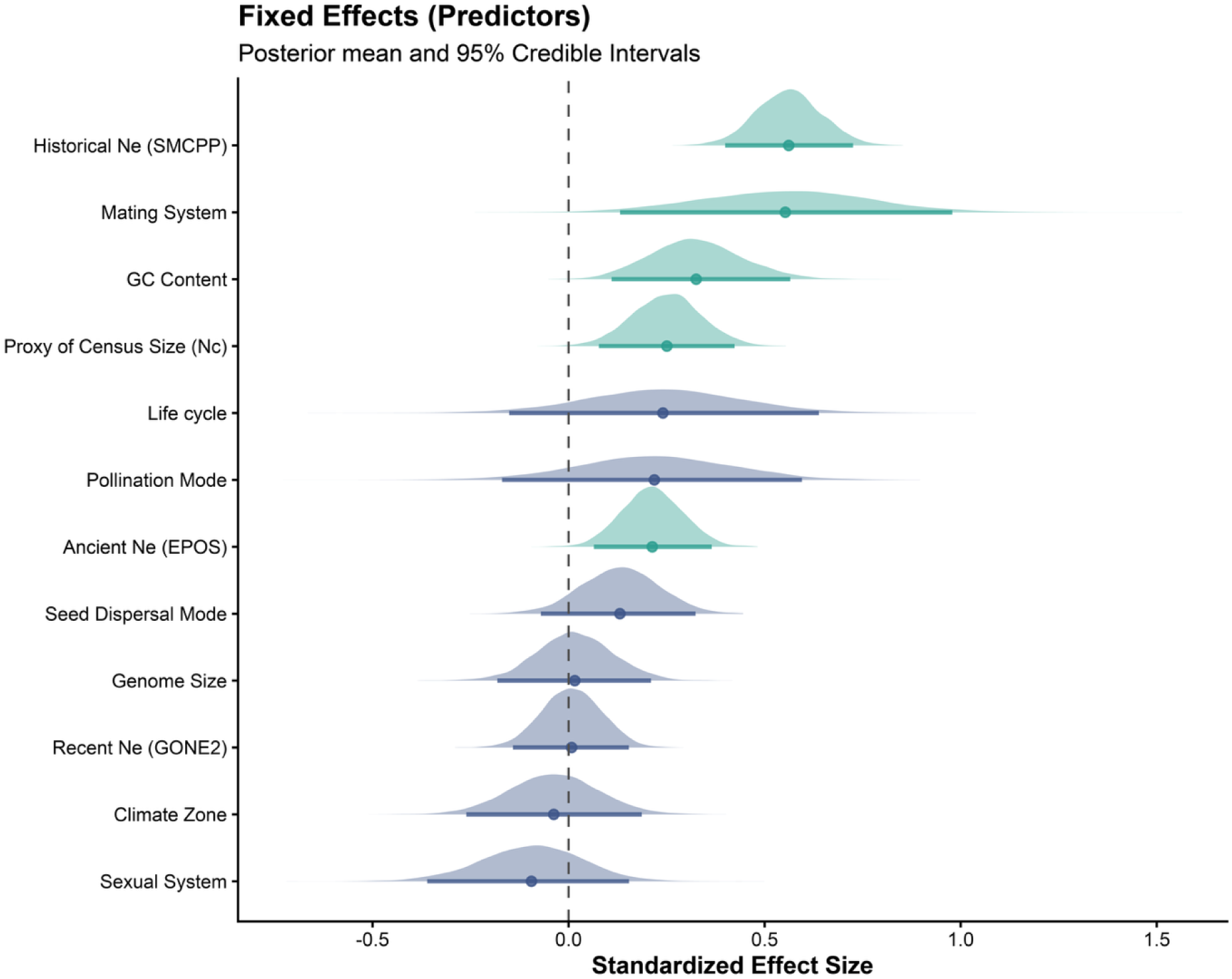
Summary of the Bayesian regression model of mixed effects on genetic diversity. Distributions of standardized effect sizes of each ecological and genetic feature. The dots represent mean values and horizontal bars represent 95% credible intervals (CI). Non-significant features (with 95% CI including zero) are shown in blue, and significant features are shown in green.

### Conservation status is mainly determined by population size

We have shown that ecological and genetic features vary in their contributions to current levels of genetic diversity. It feels then natural to ask which of those features contribute most to the species conservation status and whether the status of DD species can be reliably predicted from those features. A factor analysis of mixed data was performed to disentangle the effect of genetic diversity on current conservation status from those of life history traits and population size (**Fig. 4A**). The first ten dimensions explained 83.6% of the total variance with 19.9% explained by dimension 1 and 13.3% explained by dimension 2 (**Fig. S7**). We noticed that most of the threatened species (79.6%) were scattered on the left side of dimension 1, to which historical 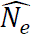, pollination mode, GC content and sexual system contributed most (**Fig. 4B**). All but five species of the non-threatened group were found on the right side of dimension 1.

**Figure 4.**
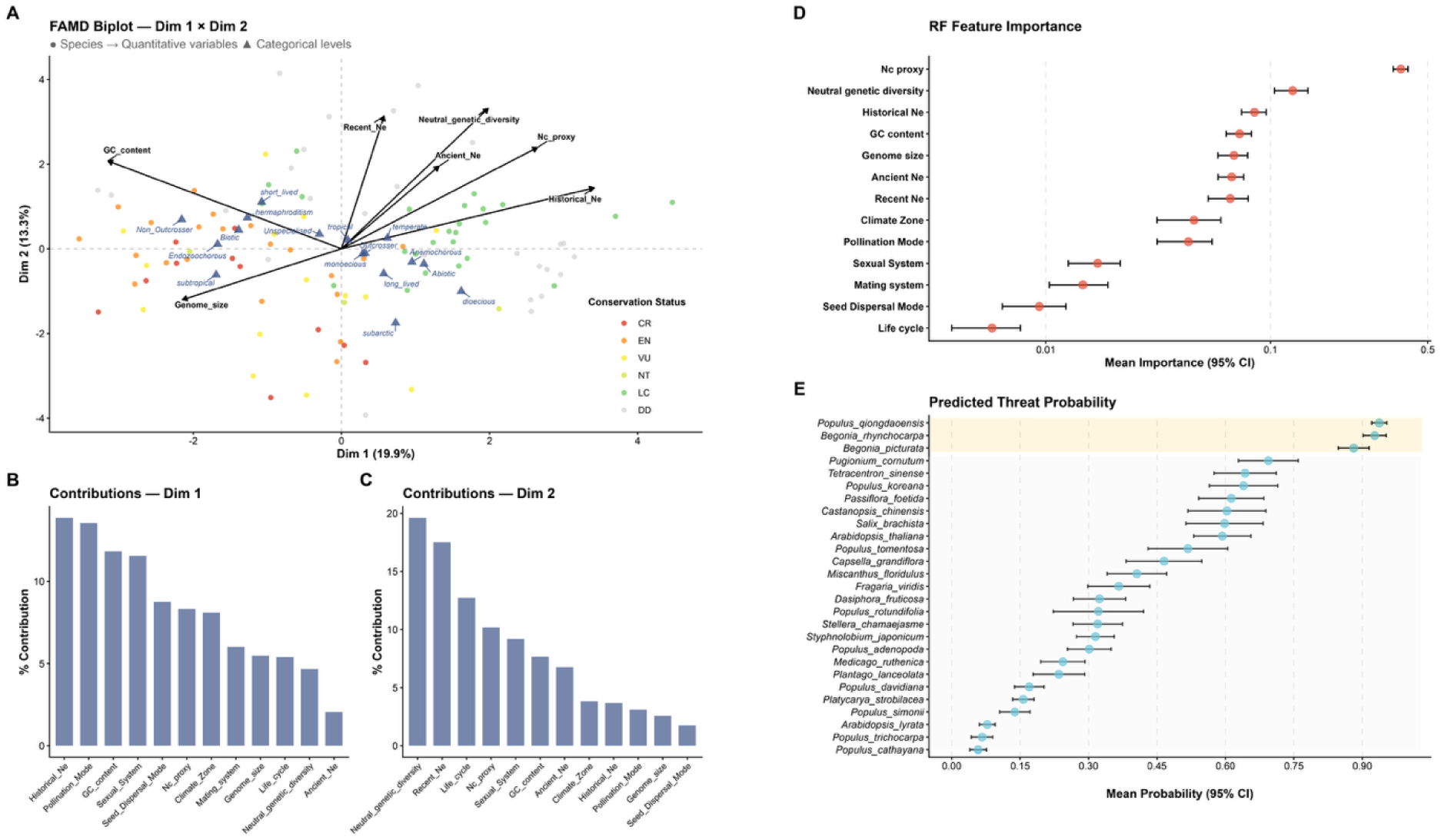
The factors affecting conservation status. (**A**) The FAMD biplot to distinguish threatened from non-threatened species. Dots represent individual species colored by their conservation status; black arrows indicate genetic variables (quantitative data), and blue triangles denote the centroids of ecological features (qualitative data). (**B and C**) The contribution of each variable to the first (Dim 1) and second dimensions (Dim 2), respectively. (**D**) Feature importance obtained from the ‘RandomForest’ model. Red points represent the mean values of importance, and error bars indicate the 95% confidence interval. (**E**) The predicted threatened probabilities for Data Deficient (DD) species based on ten random iterations. Blue points represent the mean values with error bars indicating the 95% confidence interval. Three ‘threatened’ species of high confidence are highlighted with yellow color.

To predict the conservation status of DD species, we trained a ‘RandomForest’ regression model with all genetic and ecological features of the threatened and non-threatened species. The ‘RandomForest’ model demonstrated robust predictive performance with both the mean AUROC and cross-validation accuracy equal to 0.98. 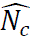 had the largest contribution (38.0%) to the conservation status prediction, followed by *π*_4_ (12.5%) and historical 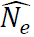 (8.5%, **Fig. 4D**). Three species in group DD (*Populus qiongdaoensis*, *Begonia rhynchocarpa*, *B. picturata*) were predicted to be threatened with high confidence across all ten iterations (95% CI of threaten probability above 0.75), six species were predicted to be non-threatened (95% CI below 0.25), while the remaining 18 could not be determined (**Fig. 4E**).

### Genetic factors can be good predictors when census size is not available

To summarize our findings, we applied a phylogenetic path analysis which revealed that census size and genetic diversity both exert direct and negative effects on species’ conservation status with the effect of census size being larger than that of genetic diversity (–0.45 versus –0.34, **Fig. 5**). Historical effective population size estimated by SMCPP, GC content, the mating system, and census size, each has an indirect effect on conservation status through their effects on genetic diversity. Moreover, historical *N_e_* also affects census size directly, which suggests species with small historical size tends to have small contemporary size. Encouragingly, an unsupervised clustering analysis based on significant genetic features (*π*_4_, SMCPP, and GC content) showed that, even when census size is not available, 69.31% of threatened and non-threatened species can be correctly assigned using other genetic and ecological factors (**Fig. 6**). However, the model had a high Type I error rate (‘genetically threatened’ but not ‘IUCN threatened’) equal to 61.22%, but a rather low Type II error rate (‘IUCN threatened’ but not ‘genetically threatened’) of 1.92%. This indicates that setting up conservation status based on three genetic features alone can have a result significantly correlated to IUCN grouping (Fisher’s exact test p-value = 1.698 × 10^-4^). However, what’s worth noting is that it could also lead to more than one third (37.03%) of ‘genetically threatened’ species being mis-classified, though, importantly, very few threatened species would be missed.

**Figure 5.**
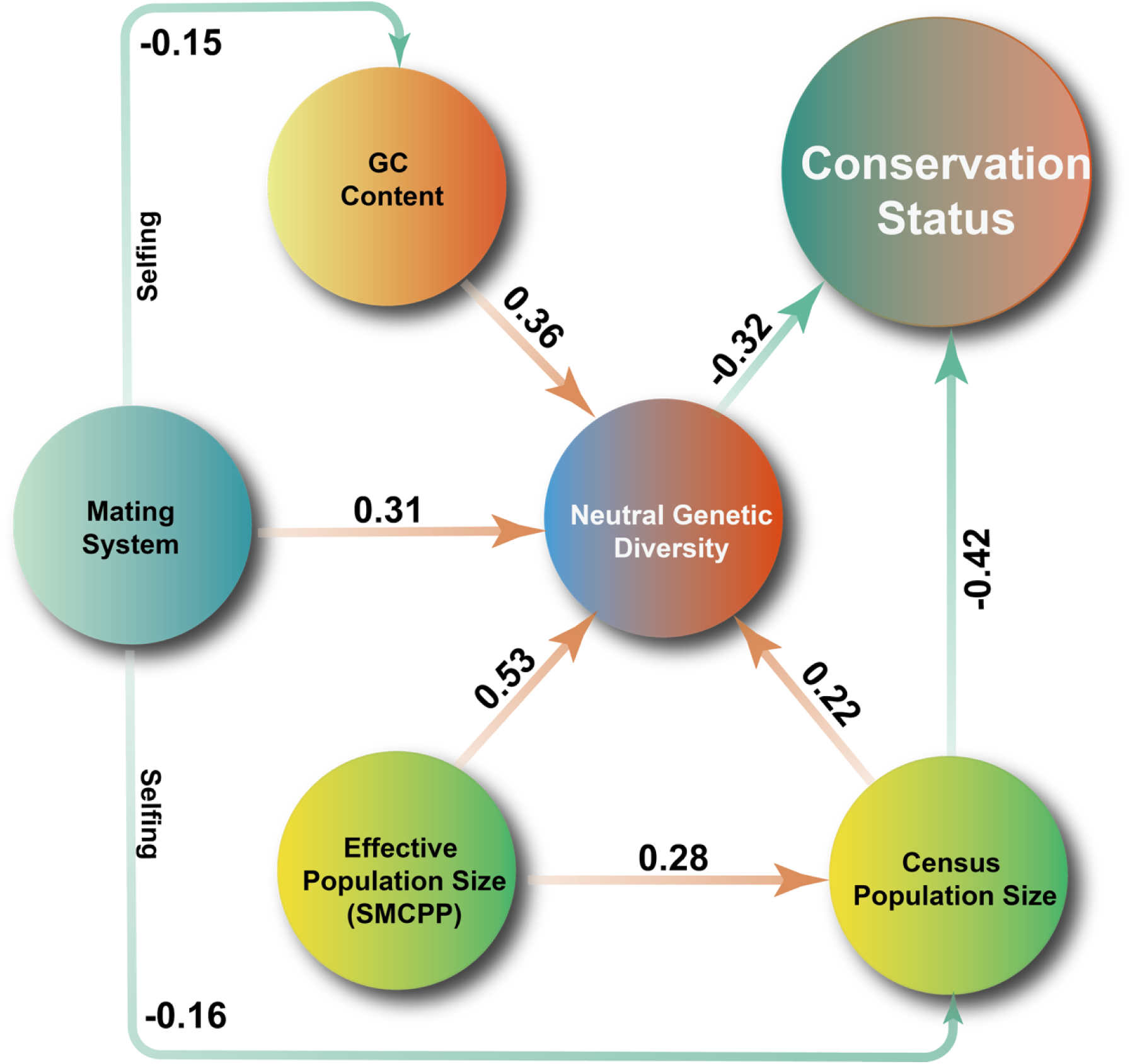
The schematic showing the major determinants of plant conservation status based on the best-fitting phylogenetic path model. Only significant effects are illustrated with values of the standardized path coefficients shown above arrows. Positive effects are highlighted in red while negative effects are in green. E.g., the negative effect of genetic diversity on conservation status means the lower the genetic diversity a species has, the more likely it will be categorized as a threatened species.

**Figure 6.**
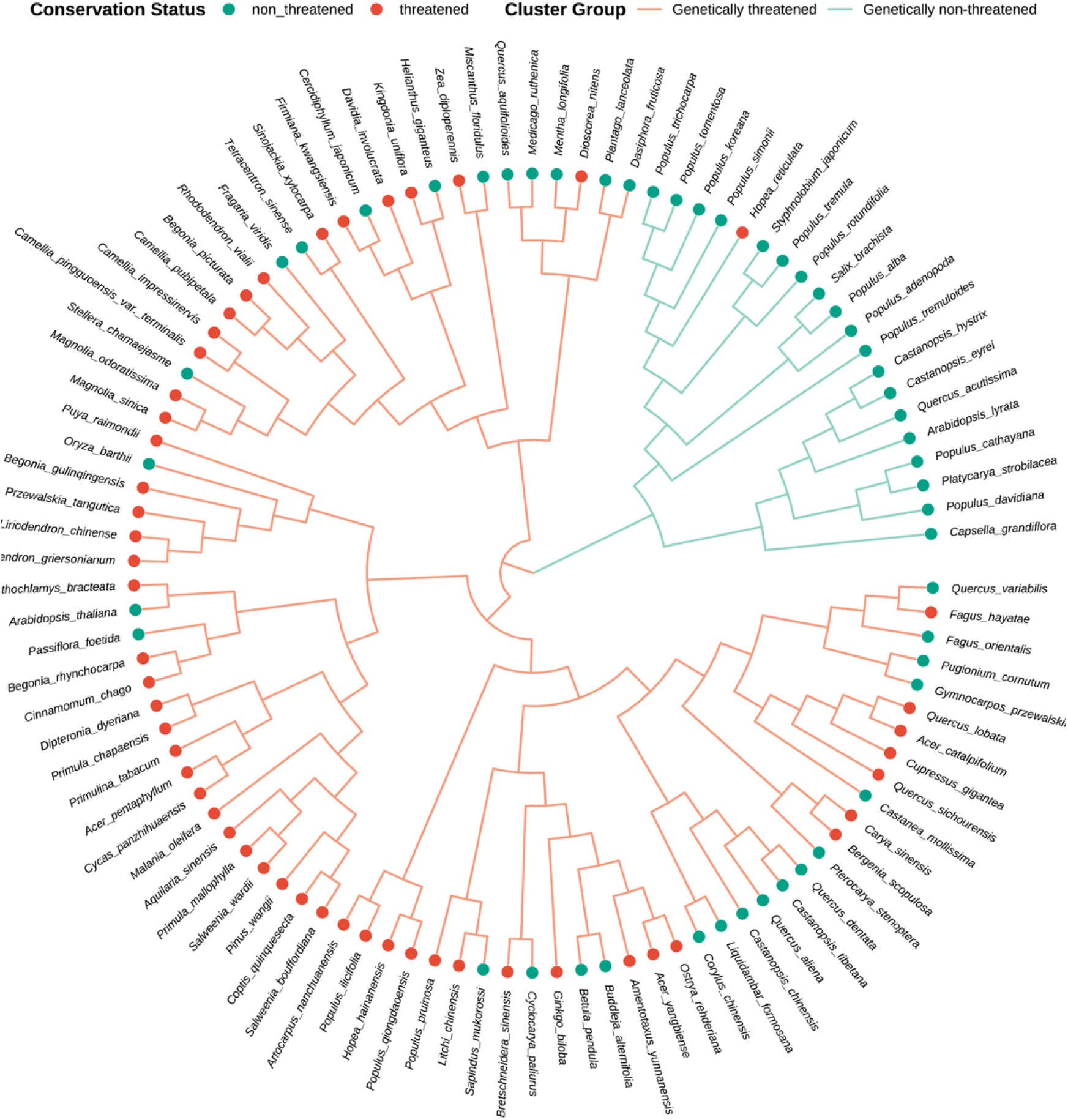
Unsupervised clustering analysis for the prediction of conservation status based on genetic features only. Circles at the periphery of the circular phylogenetic tree denote conservation status, with branch colors highlighting genetically threatened (red) and non-threatened (blue) groups. By classifying three Data Deficient (DD) species (*Populus qiongdaoensis*, *Begonia rhynchocarpa*, and *B. picturata*) as threatened and the remaining 24 as non-threatened, the clustering incorporates a total of 52 threatened and 49 non-threatened species.

## Discussion

In the present study we have used extensive genomic data in 101 species of varying census size and life history traits to assess the usefulness of different demographic and genetic parameters for the classification of species according to their conservation status. We show that threatened species have smaller *N_c_* and *N_e_*, independently of the time interval considered, lower nucleotide diversity and lower selection efficacy against deleterious mutations. As in the case of placental mammals (*5*), the ratio of historical effective population size to contemporary census population size was larger in threatened species than in non-threatened ones. Though census size can be sufficient when available (*5*), we also found that genetic diversity and historical effective population size were good predictors of the conservation status.

### Genetic diversity and demographics

There was a 100-fold difference between the lowest and highest estimates of neutral nucleotide diversity among the 101 species. This is considerably more than the 16.4-fold variation observed in a recent survey of nucleotide diversity among songbirds from the Himalayan–Hengduan region but is congruent with the variation observed by Chen *et al*. (2017) among plant species (*15, 16*). One of the main causes of this striking variation is the inclusion in both the current dataset and in Chen *et al*. (2017) dataset of species with different reproduction systems and highly variable longevity (*15*). Both reproduction systems and longevity have been shown to strongly impact nucleotide diversity in both plants and animals (*15, 17, 18*). In the present study, considering species with extremely small current census sizes also contributed to the large span in nucleotide diversity. Aside from these life-history factors and current census size, the main factor associated to the current neutral nucleotide diversity was the historical effective population size. The same conclusion was drawn by Liu *et al*. (2026) in their study of Himalayan songbirds (*16*). More specifically, they noted that nucleotide diversity “showed no correlation with recent population dynamics, current population size […]. Instead, historical demography strongly predicted genetic diversity, with ancestral population size during the late Pleistocene emerging as the sole correlate”. This may be less surprising than it seems. First, nucleotide diversity is primarily influenced by common alleles and these alleles tend to be old. For instance, in humans a neutral allele at a frequency of 0.05 arise around 200,000 years ago and a neutral allele at a frequency of 0.33 at around 700,000 years ago (*19*). The average estimated age of a neutral allele is ∼390,000 years, so roughly between the divergence of hominid lineages and the origin of modern humans. Hence, the fate and current frequency of these neutral alleles will be primarily impacted by what occurred over these time periods and the current genetic diversity reflects the demographic changes that occurred over these historical periods. Second, the historical period, as defined in the present study ([200, 10^5^] generations), is also the period where most climatic events that affected population size and structure and consequently the current genetic diversity, such as the Last Glacial Maximum (ca 20,000 ya), took place. We note, however, that in our study, the current genetic diversity was not uncoupled from the current census size. It may be that species that currently have a small census size are more likely to have experienced fluctuations in size in the past and therefore also tend to have a smaller historical *N_e_* (see below). Liu *et al*. (2026) considered only songbirds and, as they themselves noted, the diversity among them in population size and life-history traits is limited and, certainly much less than it is in our dataset (*16*). Also, they did not estimate population size but instead used abundance estimates and range size as proxies. In our study we had direct estimates of population size for 44 species, and indirect estimates of population size for the remaining species. The availability of the 44 species with direct estimates allowed us to check indirect estimates. Germain *et al*. (2023) considered 263 avian species, including Passerine and non-Passerine species and estimated past changes in *N_e_* with PSMC, so roughly from 30,000 years to 1 Mya (*20*). They distinguished 7 classes of demographic histories. However, in contrast to our study, the conservation status of the species (threatened/near threatened) was not associated to classes of demographic histories. Nonetheless, some phenotypic traits, e.g., larger clutch sizes, were associated, albeit weakly, with increasing *N*_e_ during climate warming. Finally, it should also be pointed out that our study, as well as other surveys of nucleotide diversity have left aside an important aspect of the data, namely population structure (*15, 16, 18*). Population structure certainly has an impact on effective population size. Under conservative migration, population structure can lead to an increase in *N_e_*, but under asymmetric gene flow the effect will be opposite (*21*).

### Conservation status, demography, and genetics

In an ideal world, demography and population genetics would be integrated into a science of population biology (*1, 22*). However, as pointed by both Lande (1988) and Lewontin (2004) this remains a formidable task and both are still very much separated (*1, 22*). So far, except perhaps in the management of captive populations, population genetics has primarily been used in conservation genetics for obtaining indirect estimates of elusive demographic parameters. The present study is no exception, and highlights the contribution of historical change in population size to the current conservation status of species. While genetic data may not permit a detailed reconstruction of past demography, coalescent modelling does allow the inference of past changes in genetic diversity, which themselves reflect environmental and demographic changes. As we saw above, today’s genetic diversity is intrinsically correlated to those past changes in effective population sizes. Thus, a correlation between the current conservation status of the species, that primarily reflects its current census size, and past changes in *N_e_*, may capture some deeper biological properties of the species. Said differently, a species that is today threatened was probably also demographically unstable in the past. Conversely, a species that was able to maintain a high effective population size over past climatic changes may also have the capacity to respond to current environmental changes (*23, 24*).

### Implications for conservation biology

The IUCN conservation status of species is based on five main demographic criteria: population decline rate, geographic range, population size, population restrictions, and quantitative analysis of extinction probability, none of which is easy to estimate. One of the main aims of the present study was to identify genomic-based proxies of demographic parameters that could be used to predict the conservation status. One of the conclusions is that, if such proxies are to be used, then estimates of current nucleotide diversity or historical effective population size are the best ones of those that were considered. Historical effective population sizes are undoubtedly very interesting and very informative on the evolutionary process. However, we share Frankham (2022) reservations on the inclusion of effective population sizes in IUCN criteria (*25*). Frankham was not referring to historical effective population size but instead to contemporary ones but his reservations apply to both types of *N_e_*. As pointed by both Frankham (2022) (“*N_e_* is far too complex for non-geneticists, as the literature is extraordinarily complex and confusing, such than even specialist evolutionary geneticists make mistakes”) and Ewens & Hössjer (2026) (“the concept of effective population size is quite complex and not in general well understood”), the inherent complexity of the concept of effective population size would not help clarify conservation goals (*25, 26*). Furthermore, a simple and basic fact, namely that *N_e_* relates to some property of the Wright-Fisher model is often ignored. So, we feel that simpler quantities such as neutral nucleotide diversity should be favored if the aim is solely to predict the conservation status of a species. Of course, if the aim is to understand how the interaction between a species biology and past climatic events shaped its present situation and aptitude to respond to environmental challenges then estimates of historical *N_e_* would be useful. Two additional good reasons to also tone down a bit the advocacy of the introduction of *N_e_* in conservation guidelines are that, (i) as pointed out by Lande (1988) extinction is essentially a demographic phenomenon, and (ii) understanding the biology of the species is key to conservation success and is also key to some direct estimation methods of contemporary effective population sizes (*1, 25*). The present study, by showing how the current conservation status could be related to past and current attributes of the species, made a significant contribution to our understanding of the biology of these species, even though additional studies are obviously needed to clarify the nature of these associations.

## Materials and Methods

### Plant samples and data collection

We compiled two datasets comprising whole genome sequencing data of 1933 individuals from 101 plant species. The first dataset includes 313 individuals from 21 endemic plant species with very narrow distribution ranges and limited numbers of mature individuals in the wild. They belong to the Plant Species with Extremely Small Populations (PSESP) program, an initiative aimed at monitoring the status of threatened plant species in Southwest China (*27*). Estimates of population sizes based on field survey are available for all species of PSESP. Fresh leaves were collected and kept at –80℃ before whole genome resequencing (WGS). DNA extraction and library preparation were performed using standard protocols by Novogene Corporation, followed by short read WGS (150 bp paired-end mode) with DNBSEQ-T7 sequencer. Six of the reference genomes used in this study were sequenced, then *de novo* assembled and annotated (see **Supplementary Texts 1.1** and **1.2**) while other reference genomes were downloaded from the public domain.

The second dataset comprises genomes of 1620 individuals from 80 plant species with distribution ranges of varying size but that are, in general, much larger than those of PSESP species except for 20 species with similar distribution ranges to those of PSESP. On average, 10–20 samples covering the whole distribution range for each species were chosen to account for available information on population genetic structure. For each species, individual genomes with significant signals of genetic admixture, i.e., with the proportion of dominant ancestral component < 0.9 in ADMIXTURE analysis, were removed. All genomic data in this second dataset were downloaded from the public domain. See **Table S1** for detail information of reference genomes and the project numbers of all downloaded data.

### SNP calling

For each species, clean short reads of genome resequencing were aligned to the reference genomes of the focal species or a closely related one using BWA-MEM (v.0.7.17) (*28*). Individual genetic variants were identified using ‘HaplotypeCaller’ in GATK (v.4.6.0) (*29*) and joint genotypes of all individuals of a given species were called using ‘GenotypeGVCFs’. To get an accurate estimate of genetic diversity and related summary statistics, both variant and invariant positions were retained in the final VCF file with the ‘--all-sites’ option (*30*). Positions of low quality were filtered out using a five-step pipeline combining GATK ‘VariantFiltration’, GenMap, PanDepth, and Bcftools. The final quality-controlled datasets, including both invariant sites and variant ones (single nucleotide polymorphisms, SNPs), were used for downstream analyses.

### Estimation of population genetic summary statistics

We calculated pairwise nucleotide diversity at four-fold degenerate sites (*π*_4_) as neutral genetic diversity, at zero-fold degenerate site (*π*_0_), and the ratio *π*_0_/*π*_4_, a measure of the efficacy of selection (*15*). Other population genetics summary statistics, including Watterson’s theta (*θ_w_*), individual heterozygosity (*H_e_*), and Tajima’s *D*, were also calculated using piawka (v.0.8.11) (*31*). For monoecious or hermaphroditic species, selfing rates were estimated using the program *roh-selfing* (*32*). Briefly, runs of homozygosity (ROHs) were first identified using the ‘*roh*’ function in Bcftools based on the SNPs dataset. Inbreeding coefficient (*F*) and Tajima’s *D* were then calculated using PLINK (v.1.9) (*33*) and VCFtools (v.0.1.16) (*34*), respectively. Finally, the selfing rate was estimated for each species using the ‘RandomForest’ model implemented in *roh-selfing* with ROH, *F* and Tajima’s *D* as the input.

### Inference of population sizes

To estimate the census population size (*N_c_*) for species for which it was not available, species distribution range was first estimated by counting the grids of size 0.1° (latitude) × 0.1° (longitude) using occurrence records retrieved from the Global Biodiversity Information Facility (GBIF). A log-log linear regression model was then established between *N_c_* and species distribution range for tree and non-tree (shrubs and herbaceous) species with known *N_c_*, respectively. Finally, *N_c_* was calculated for the remaining species for which direct estimates of *N_c_* were not available, based on the linear regression models established in the previous step.

Effective population size (*N_e_*) was estimated over three different time intervals: [10^5^,10^7^], [200, 10^5^], and [1, 200] generations (referred to as ‘ancient’, ‘historical’, and ‘recent’ *N_e_*, respectively). Firstly, the SFS-based method EPOS (v.1.7.2), was used to estimate *N_e_* changes in time interval [10^5^,10^7^] based on unfolded SFS (option ‘-U’) (*12*). The unfolded SFS was constructed using EST-SFS (v.2.0.4) for non-missing, four-fold degenerate sites with two outgroups (*35*). Secondly, SMCPP (v.1.15.2), a method based on the distribution of heterozygosity along the genome, was used to capture *N_e_* changes in time interval [200, 10^5^] with default settings (‘--timepoints 2e2 1e5’) (*13*). Repetitive regions were excluded and the individual with the highest sequencing coverage was chosen as the “distinguished” sample (via the option ‘-d’). Thirdly, a linkage disequilibrium based method, GONE2, was applied to estimate *N_e_* changes in time interval [1, 200] with the option ‘-x’ accounting for population structure (*14*). The mutation rate *μ* was set to 1.28 × 10^−8^ per base pair per generation for all species (*36*) for the following reasons: 1) the mutation rate and generation time are not available for most species and are difficult to estimate, especially for perennial species with overlapping generations; 2) A recent study in animals showed that the mutation rate per base pair per generation is very uniform (*37*).

Finally, we computed the harmonic mean (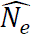) over the three intervals [10^5^,10^7^], [200, 10^5^], and [1, 200] generations using the following equation:

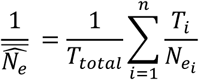

Where *T_ii_* is the length of time tile (*i*) in unit of generation; *N_ei_* is the estimate of effective population size over that time tile; *n* is the number of different *N_e_* during the total time interval and *T_total_* is the total length of time tiles in which *N*_e_ estimates is obtained.

### Species features and conservation status

All species were characterized by ecological features, which include life cycle (short-lived or long-lived), growth form (herbaceous, shrubby, or woody), sexual system (monoecious, dioecious, or hermaphroditic), seed dispersal (anemochory, endozoochory, or unspecialized), pollination mode (biotic or abiotic) (*38, 39*), and climate zone (subarctic, temperate, subtropical, or tropical) that were obtained from Plants of the World Online (https://powo.science.kew.org/). All dioecious species together with monoecious or hermaphroditic species with selfing rate < 0.2 were regarded as outcrossing mating system while the rest were considered as non-outcrossing (partially or strictly selfing). Genomic features were also calculated for all species, including genome size (in Gigabase), repetitive content, and GC content. Species conservation status in IUCN Red List of Threatened Species (https://www.iucnredlist.org/, last visit on November 16, 2024) or the Information System of Chinese Rare and Endangered Plants (https://www.iplant.cn/bhzw/; last accessed October 26, 2024) were used to group all 101 species into three categories: 1) 49 ‘threatened’ species that are classified as Critically Endangered (CR, n=11), Endangered (EN, n=21), Vulnerable (VU, n=14), or Near Threatened (NT, n=3); 2) 25 ‘non-threatened’ species that are classified as Least Concern (LC, n=25); and 3) 27 ‘data deficient’ (DD) species since they have not been included in any conservation programs by now and their conservation status still needs to be identified.

### Identifying determinants of genetic diversity

We first performed pairwise correlation tests on 23 ecological and genetic features to remove multicollinearity. In total 12 features were retained with a correlation coefficient *r* < 0.40 for quantitative variables and *r* < 0.55 for qualitative traits, including six ecological features (mating system, seed dispersal, pollination, sexual system, life cycle, and the climatic zone of the distribution range), and six genetic features (three estimates of 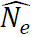, 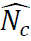, genome size and GC content).

We then regressed neutral nucleotide diversity (*π*_4_) on these 12 features using Phylogenetic Location-Scale Models (PLSMs) implemented in a Bayesian framework in the R package ‘brms’ (*40*). All 12 features were considered as fixed effects and *π*_4_ and all numeric features were first normalized using the R package ‘bestNormalize’ (*41*). A covariance-correlation matrix was derived from the phylogenetic tree using the R package ‘rtrees’ (*42*) and used to estimate the phylogenetic signal (*λ*) as a random effect in the model (*43*). Each of four independent MCMC chains was run for 4,000 iterations (2,000 burn-in) using default parameters.

### Conservation assessment and predictive modeling

To identify the main factors affecting conservation status, we applied a Factor Analysis of Mixed Data (FAMD) using the ‘factoextra’ R package (*44*) for all species and the 12 features described in the “*Identifying determinants of genetic diversity*” section. Furthermore, we predicted the conservation status of the 27 Data Deficient species using a ‘RandomForest’ classifier implemented in the Python library ‘scikit-learn’ (*45*). The 74 species with known conservation status and 12 features were randomly partitioned into a training (75%) and a test (25%) set. Hyperparameters were optimized via cross-validation to refine the decision tree structure, and model performance was evaluated using the Area Under the Receiver Operating Characteristic Curve (AUROC). Feature importance was calculated by averaging the predictive contribution of each variable across ten randomized iterations. Given that *N_c_* is the main criterion used by IUCN and other conservation agencies, we also performed an unsupervised hierarchical clustering analysis on all species with all features but *N_c_* and conservation status. The optimal number of clusters was determined using the ‘fviz_nbclust’ function in the R package ‘factoextra’ (*44*). Finally, we conducted a phylogenetic path analysis using the R package ‘phylopath’ (*46*) to summarize the effects of historical *N_e_* estimated by SMCPP, *N_c_*, mating system, genome size and GC content on genetic diversity and conservation status.

Please refer to **Supplementary Materials and Methods** for more detailed descriptions of all methods described above.

## Data availability

All data generated for the study are deposited in the China National Center for Bioinformation-National Genomics Data Center (https://bigd.big.ac.cn/) website with accession no. CRA029742 and CRA030844 (reviewer links: https://ngdc.cncb.ac.cn/gsa/s/ZZ5s7CP7 and https://ngdc.cncb.ac.cn/gsa/s/7w5c8dft). Data for *de novo* assembled genomes have been deposited with a BioProject number PRJCA044520 (a reviewer link: https://ngdc.cncb.ac.cn/gwh/Assembly/reviewersPage/yLfwJvRULUJSIxARBjjqbFSinqoBzitaMwXQGIX GTSnnQGStkXRjDvrfuPQnmRfW). All genomic data will be released and public upon accepted.

## Code availability

All code used for the analyses are available (for reviewer link: https://www.protocols.io/blind/1E09D7010CAA11F1BBE80A58A9FEAC02).

## Acknowledgments

This research was financially supported by the Key R&D Program of Zhejiang (2024C03244) and the National Natural Science Foundation of China (32371689) to Chen Jun; the Science and Technology Projects of the Xizang Autonomous Region, China (XZ202402ZD0005) and the China Conservation Collaboration Initiative to Pan Li. The computations were enabled by resources provided by the National Academic Infrastructure for Supercomputing in Sweden (NAISS) at UPPMAX (Projects NAISS 2024/6-9 and UPPMAX 2025/2-230), funded by the Swedish Research Council through grant agreement no. 2023-03904 to Martin Lascoux. The paper was in part written during a one-year visit of the first author to Uppsala University. This visit was financed by Zhejiang University.

We thank Weibang Sun and the Molecular Biology Experiment Center at the Germplasm Bank of Wild Species for providing plant samples in this study; Meizhen Wang for her assistance with sample collection; Ruirui Fu, Jing Wu, Junjie Wu and Yuting Jiang for their valuable discussions and insights; and Yang Liu for providing photographs of our focal plant species.

## Author Contributions

J.C. and M.L. designed and managed the project; T.T.Z, Y.X.Z., S.N.Z., M.M.Z., Y.L., P.L., W.B.S. collected plant materials; T.T.Z. analyzed the data; T.T.Z., M.L., and J.C. wrote the manuscript.

## Classification

major: Biological Science; minor: Evolution

