## Supplementary material for "The contribution of recent and historical demographic histories to genomic diversity and conservation status in plant species"

PNAS

**This PDF file includes:**

Supporting text
Figures S1 to S16
SI References

### Supporting Information Text

#### 1. Materials and Methods

##### *1.1 Genome sequencing and assembly*

In this study, we sequenced and *de novo* assembled genomes of six species, *Amentotaxus yunnanensis*, *Begonia gulinqingensis*, *Firmiana kwangsiensis*, *Primula chapaensis*, *Primula mallophylla*, and *Salweenia bouffordiana*. For each species, fresh leaves were collected and frozen in -80℃ before being sent to Novogene Corporation (Beijing) for sequencing. Whole genomic DNA was extracted using the QIAGEN Genomic DNA extraction kit (QIAGEN, Hilden, Germany) following the manufacturer's protocol and PacBio libraries were prepared using the SMRTbell Express Template Prep Kit 2.0 (Pacific Biosciences, USA) following PacBio's standard protocol with an insert size of 15 kb. PacBio HiFi reads (~20× coverage) were produced on the Pacific Biosciences Sequel II platform in circular consensus sequencing (CCS) mode.

Sequencing adaptors were removed from raw HiFi reads using HiFiAdapterFilt (*1*) before *de novo* assembly. Clean reads were converted to FASTQ format using 'bam2fastq' module implemented in PacBio BAM toolkit (https://github.com/PacificBiosciences/pbtk, last accessed 2024.03.12) and *de novo* assembled into contigs using HiFiasm (v.0.19.8) (*2, 3*) following default parameters and the authors' recommendations. Please refer to **Table S2** for all summary statistics for genome sequencing and assembly.

##### *1.2 Gene annotations*

Firstly, full length transcriptomes were sequenced and assembled in *Firmiana kwangsiensis* and *Salweenia bouffordiana* and were used for gene annotations. Protein homolog sequences were available in closely related species for the four other species with, and gene annotations were performed based on protein homolog sequences. Total RNA was extracted from fresh leaves using Trizol reagent (Invitrogen, CA, USA). PacBio full-length isoform libraries (Iso-seq, Pacific Biosciences, CA, USA) were prepared using Sequel Iso-seq Express Template Prep (Pacific Bioscience, USA) according to the manufacturer's protocol and then sequenced on the PacBio Sequel platform (PacBio, USA). Raw Iso-Seq reads were processed using the IsoSeq pipeline (https://isoseq.how/) to obtain polished consensus sequences. Specifically, circular consensus sequences (CCSs) were constructed from the raw subreads by using ccs (v.6.0.0) (https://github.com/PacificBiosciences/pbbioconda) with the following parameters "–skip-polish–min-passes 4–min-length 200–min-rq 0.99". The primer sequences in the long CCS reads were removed by using lima (v.2.0.0) with the parameter "–isoseq". The full-length, nonchimeric (FLNC) cDNA reads were identified by the refine subcommand of isoseq (v.3.4.0) with the following parameters: "–require-polya–min-polya-length 20". Then, we used the cluster and polish subcommand to cluster FLNC reads and generated polished transcripts.

Secondly, repetitive elements were identified in genome assembly with HiTE (v.3.1.1) (*4*). Protein coding genes were then predicted from soft-masked genomes using BRAKER3 pipeline (https://github.com/Gaius-Augustus/BRAKER?tab=readme-ov-file#braker-with-short-and-long-read-rna-seq-and-protein-data) (*5*). GeneMark-ETP was trained using both species' polished transcripts and orthologous protein sequences from 416 plant species (obtained from OrthoDB 12, https://bioinf.uni-greifswald.de/bioinf/partitioned_odb12/) (*6*) in order to generate a set of high confident genes, which was later used to train AUGUSTUS (*7*). The predictions of GeneMark-ETP and AUGUSTUS were combined using TSEBRA to generate the final protein coding gene models.

For gene annotations of the four other reference genomes (*Begonia gulinqingensis*, *Primula chapaensis, P. mallophylla,* and *Amentotaxus yunnanensis*), Lifton (v. 1.0.1) (*8*) was used to predict protein coding genes based on homolog genes from genomes of related species. Particularly, we used *Begonia masoniana* (*9*) homolog genes to annotate *B. gulinqingensis* genome, used *Primula veris* (*10*) homolog genes to annotate *P. chapaensis*, and *P. mallophylla* genomes and used *Taxus wallichiana* (*11*) homolog genes to annotate *Amentotaxus yunnanensis* genome.

#### *1.3 Genome assembly evaluation*

To evaluate the completeness of genome assembly and gene annotations for the six species, BUSCO (v. 5.4.5) (*6*) scores were calculated based on 1,614 protein sequences in embryophyta odb10 database. Additionally, PanDepth (v. 2.25) (*12*) was applied to evaluate genome completeness based on mapping coverage with PacBio HiFi reads and DNBSEQ-T7 short reads.

#### *1.4 Population resequencing*

We generated population genomic resequencing data for 21 species that belong to the Plant Species with Extremely Small Populations (PSESP) program in Southwest China. Fresh leaves were collected for two species (*Cycas panzhihuaensis* and *Cinnamomum chago*, one site per species) while silica gel–dried leaves were used for the remaining 19 species (one to five sites per species; **Table S3**). Ten to twenty individuals were randomly selected for each species, and in total 313 samples were used for whole-genome resequencing. Genomic DNA was extracted using the QIAGEN Genomic DNA extraction kit (QIAGEN, Hilden, Germany) following the manufacturer's protocol. A total of 0.2 μg DNA per sample was used as input material for the short read library preparation using the NEBNext® Ultra™ II DNA Library Prep Kit (New England Biolabs, Ipswich, MA, USA). Genomic DNA was first fragmented by sonication to reach a size of 350 bp. Next, the DNA fragments underwent end polishing, A-tailing, and ligation with the full-length adapter for sequencing, followed by PCR amplification. The PCR products were then purified using the AMPure XP system (Beverly, USA). The 5'-end of each library was phosphorylated and cyclized. Then, loop amplification was performed to generate DNA nanoballs. Finally, these DNA nanoballs were loaded into a flow cell with DNBSEQ-T7 for sequencing.

#### *1.5 Short read alignment and SNP calling*

For each species, sequencing adapters were trimmed from raw reads, and reads containing low-quality bases (<5) or ambiguous nucleotides ('N's) were removed using Fastp (v.0.23.1) (*13*). Clean reads were then aligned to the reference genomes of a focal or closely related species using BWA-MEM (v.0.7.17) (*14*). Reads of PCR duplication were identified and removed using streammd (v.4.3.0) (*15*) and all aligned reads were sorted by genomic coordinate with Sambamba (v.1.0.1) (*16*). Individual genetic variant calling was performed using 'HaplotypeCaller' in GATK (v.4.6.0) (*17*) and a genomic variant call format (GVCF) file was generated for each individual. All GVCFs files of a given species were combined and the joint genotypes of all individuals were called using 'GenotypeGVCFs' in GATK. Both variant and invariant positions were retained in the final VCF file for each species using the '--all-sites' option in order to estimate the genomic diversity and related summary statistics accurately (*18*).

A quality control pipeline was conducted on VCF files with the following steps: 1) the genetic variants were first filtered using GATK's 'VariantFiltration' module with the following thresholds 'MQ < 30.0, MQRankSum < –20.0, QD < 2.0, QUAL < 30.0, SOR > 3.0, FS > 60.0, and ReadPosRankSum < –8.0'; 2) For all positions along the entire genome, GenMap (v.1.3.0) (*19*) was used to calculate the mappability, which is a measure of how unique or repetitive a genomic region is. Sites with mappability equal to 1 (i.e. completely unique) were retained using Bcftools (v.1.20) (*20*) while sites that contained missingness or were located in repetitive regions (based on gff file inferred by HiTE with default settings) were removed; 3) PanDepth was employed to calculate the sequencing depth for each species and we only kept the sites that had mean depth of more than 10; 4) We filtered out variants that deviated significantly from Hardy–Weinberg equilibrium (HWE ≤ 0.05) using Bcftools (**Fig. S8**). The final quality-controlled datasets, including both invariant sites and variant ones (single nucleotide polymorphisms, SNPs), were used for downstream analyses (**Table S4**).

#### *1.6 Calculating genomic properties*

We calculated the genome size (Gb) and GC content (%) using seqkit2 (*21*) for 83 species for which reference genomes are available either in the public domain or sequenced in this study. The repetitive content proportions (%) were estimated with HiTE with default options, which systematically identified and masked transposable elements across each species genome. Moreover, for 18 species without genomes, 14 reference genomes of closely related species within the same genera of the focus species were used to estimate genome size, GC content, and repetitive content (See list in **Table S5**). Considering the possible variation among species even within same genus, we also performed all analyses related to these three genomic properties using 83 species (see section **2.6**).

#### *1.7 Census size estimation*

Firstly, the distribution range was estimated for each species based on occurrence records from the Global Biodiversity Information Facility (GBIF; last accessed 24 March 2026). Only occurrences recorded between

1960 and 2025 were used. We also used the information from the Plants of the World Online (POWO) database (https://powo.science.kew.org/) to remove occurrence records that do not belong to the species native distribution range. The R package 'CoordinateCleaner' (v. 3.0.1) (*22*) was used to remove problematic records including duplicates, records of erroneous coordinates (e.g., oceanic coordinates, city, arboretum, or missing, zero, or inverted coordinates), fossil or living collection records, and so on. The cleaned occurrence records were projected into grids of size 0.1° (latitude) × 0.1° (longitude) (approximately 11.1 × 11.1 $km^2$). The total number of unique occupied grid cells was counted for each species and used as species distribution size. Secondly, a log-log linear regression model was established between census size ($N_c$) and species distribution size for 44 species for which Nc was available and for tree and herbaceous/shrubby species, respectively. In total, $N_c$ from field survey was available for 44 species (provided by the Kunming Institute of Botany, KIB and from IUCN RedList; **Table S6**) with the population genetics data of 21 species also included in the dataset of 101 species in this study. Both distribution size and $N_c$ were log10-transformed and the coefficient and intercept of the linear regression model was estimated using function 'lm' in R package 'stats' (v.4.4.3) for trees and herbaceous/shrubby species, separately. Finally, for the remaining 80 species in this study, $N_c$ was estimated using the log-log linear regression model (depending on whether they are tree or herbaceous/shrubby species) and their distribution sizes with the 'predict' function from the 'stats' R package.

Notably, for *Quercus lobata*, an oak species endemic to California (in interior valleys and foothills from Siskiyou to San Diego counties), a recent burst of GBIF records was observed from 2017 to 2025 by comparing with IUCN records (https://www.iucnredlist.org/species/61983021/61983023), which led to an overestimate of $N_c$ of around 20 million. Such increase of GBIF records could be caused by scientific survey for conservation purpose. To keep data consistency among species, we therefore only used *Q. lobata* GBIF records from 1995 to 2016.

#### *1.8 Effective population size estimation and regression*

*EPOS:* We utilized EPOS (v. 1.7.2) (*23*) to estimate ancient effective population size ($N_e$). First, easySFS.py (https://github.com/isaacovercast/easySFS) was used to generate a one-dimensional unfolded site frequency spectrum (SFS). This was based on a derived-SNP dataset restricted to four-fold degenerate sites (treated as nearly neutral) without any missing sites. Ancestral states for each site were inferred using EST-SFS (*24*) with two outgroup species. As EPOS requires the number of both monomorphic and polymorphic sites, we did not apply any minor allele frequency filtering to the dataset. We ran EPOS with the mutation rate set to $1.28 \times 10^{-8}$ (option '-u'), providing the neutral unfolded SFS (option '-U') alongside the total number of monomorphic and polymorphic sites (option '-l') as input. Finally, we restricted our analysis to the resulting $N_e$ curve located within the generation time interval of [$10^5$, $10^7$].

*SMCPP*: VCF files were converted to SMC format using 'smcpp vcf2smc' with SNPs located in repetitive regions excluded. For each species, one individual with the highest sequencing coverage was used as a 'distinguished' sample (via the option '-d') using SMCPP (v. 1.15.5) (*25*). Historical $N_e$ was estimated using ‘smcpp estimate’ with default settings (‘--timepoints 2e2 1e5’), which set unit of time $T$ to generation $N_e$ spanning from 200 to $10^5$ before present (refer to Supplementary PDF-File “SMCPP-Ne-curve”).

*GONE2*: Neither EPOS nor SMCPP was designed to capture very recent changes in $N_e$. In contrast, GONE2 leverages the spectrum of linkage disequilibrium (LD) between pairs of loci in a population to estimate demographic history over the past two hundred generations and population structure was considered using the “-x” option. In principle, inference of $N_e$ with GONE2 (v.1.0.2) (*26*) requires a genetic map. However, a genetic map is unavailable for most of non-model plant species in our study. Instead, we estimated a mean population recombination rate ρ[1/bp] across the genome with the LDhat software (v.2.1) (https://github.com/auton1/LDhat). Recombination rate, r[cM/Mb], was then obtained by using the relationship r = $\mu \times \rho/\theta$ for each species, which was scaled to the unit of cM/Mb.

Next, we also calculated a statistic $N_e\_decline$ that describes the maximum demographic fluctuation based on $N_e$ changes in the historical $T$ interval:

$$N_e\_decline = 1 - \frac{N_e^{MIN}}{N_e^{MAX}}$$

where $N_e^{MIN}$ and $N_e^{MAX}$ correspond to the minimum and maximum effective population sizes, respectively, of the steepest continuous decline event across time tiles of each species.

Finally, we fitted the $N_e$ trajectories using the generalized additive model with cubic regression splines for all threatened and non-threatened species and for three different time scales, respectively. All curves with 95% confidence intervals were visualized using the ‘geom_smooth’ function in the R package “ggplot2” (v.4.0.2). Notably, only in this analysis, 49 threatened species plus three predicted threaten species in group DD (*Populus qiongdaoensis*, *Begonia rhynchocarpa*, *B. picturata*, based on results of ‘RandomForest’ classifier) were used to fit the ‘threatened’ cubic regression spline while the remaining 52 species in non-threatened and DD groups were used to fit the ‘non-threatened’ cubic regression spline.

#### *1.9 Phylogenetic regression analysis*

Before evaluating which features contribute most to genetic diversity, we first carried out a pairwise correlation test to remove multicollinearity between the 23 ecological and genetic features using ‘cor.mtest’ function (for quantitative traits) and ‘assocstats’ function (for qualitative traits) in the R packages ‘corrplot’ (*27*) and ‘vcd’ (*28*), respectively (see **Table S1** for all these features). In total 12 features were retained with a correlation coefficient $r < 0.40$ for quantitative variables and $r < 0.55$ for qualitative traits, including six ecological features (mating system, seed dispersal, pollination, sexual system, life cycle, and the climatic zone of the distribution range), and six genetic features (three estimates of $\widehat{N_e}$, $\widehat{N_c}$, genome size and GC content). We then conducted the Phylogenetic Location-Scale Models (PLSMs) by regressing the neutral nucleotide diversity ($\pi_4$) on these 12 features using the R package ‘brms’ (*29*). $\pi_4$ and all numeric features were first normalized using the R package ‘bestNormalize’ (*30*). All 12 features were taken as fixed effects. The phylogenetic tree of 101 species was transformed into a covariance-correlation matrix using the R package 'rtrees' (*31*) and used to estimate the phylogenetic signal ($\lambda$) as a random effect in the model (*32*). Considering the influence of uncertainty in branch lengths on regression analyses, the total branch length from the most recent common ancestor to the species tips were normalized to one, following the recommendations of Symonds *et al* (2014, p104-130) (*33*). A phylogenetic signal ($\lambda$) approaching 1, indicates that a correlation between species is expected under a Brownian model of evolution (i.e., strong phylogenetic dependency) while A phylogenetic signal, $\lambda$, closer to zero indicates that the species are less correlated and less dependent on the phylogenetic structure. To estimate the effects and significance for all features in PLSM, four independent MCMC chains were run for 4,000 iterations with 2,000 burn-in using default settings.

#### *1.10 Phylogenetic path analysis*

To summarize the effects of major features on the conservation status, we conducted a phylogenetic confirmatory path analysis using the R package 'phylopath' (v. 1.3.1) (*34, 35*). Briefly, a generalized least-square model (PGLS) implemented in the R package 'phylolm' is used to take account for the phylogenetic effect (*36*). Then an acyclic graph is established by finding and ordering the significant causal effects based on the D-separation test implemented in the R package 'ggm' (https://github.com/cran/ggm). For this analysis, we included mating system (outcrossing, partly or strictly selfing), GC content, historical effective population size (estimated by SMCPP), neutral genetic diversity, and census population size, which had significant influence on either genetic diversity or conservation status in previous analyses.

We first defined a baseline structural model in which four predictors, namely, $\widehat{N_e}$, $\widehat{N_c}$ , GC and the mating system that had a significant effect on $\pi_4$ in previous PLSM analysis also had an independently influence on $\pi_4$. We then used this model as the foundational structure to sequentially test the following four models: 1) Model 1, conservation status is only determined by $\widehat{N_c}$ and $\pi_4$; 2) Model 2, add an effect of the mating system on GC content to Model 1 (*37, 38*); 3) Model 3, add an effect of the mating system on $\widehat{N_c}$ to Model 2; 4) Model 4: add an effect of $\widehat{N_e}$ on $\widehat{N_c}$ to Model 3 considering the influence of historical demographic fluctuations on current census size (*39*). **Fig. S9** provides a schematic of each of the four structural models.

Each effect within the model was formalized as a directed acyclic graph (DAG), which encodes conditional independence ("D-separation") statements among variables. A conditional independence statement in D-separation test dictates that two variables should be statistically uncorrelated once appropriate confounding variables are controlled for (e.g., if A affects B solely through mediator C, A and B should be independent when conditioning on C). Violations of these expected independencies indicate that the model omitted one or more necessary causal pathways. The significance of independence in D-separation test was calculated using Fisher's C-statistic: a low C-statistic with a non-significant *p*-value indicates an acceptable model, whereas a significant value ($P < 0.05$) suggests that at least one assumption of conditional independence was violated. All effects within each model were assessed using D-separation tests that also accounted for phylogenetic nonindependence (*40*).

To compare models, we first computed the C-statistic information criterion corrected for small sample sizes (CICc) that balances model complexity with explanatory power. ΔCICc was then generated by subtracting the lowest CICc value from the model's original CICc value. Models were ranked by ascending ΔCICc values with ΔCICc of the top ranked model equal to zero. The top ranked model is selected as the best model if ΔCICc of the second top ranked model is larger than two. Otherwise, all models with ΔCICc smaller than two will be as taken as acceptable models and model coefficient values will be averaged based on 'Akaike weight', $\omega_i$, which can be calculated as:

$$\omega_i = \frac{\exp(\Delta_i/2)}{\sum_{r=1}^{R} \exp(\Delta_r/2)}$$

where $\Delta_i$ is the ΔCICc of the $i$-th model and $R$ is the number of models.

Finally, we used standardized path coefficients to evaluate the effect of each variable within the best model. For multiple acceptable models, a mean coefficient weighted by $\omega$ was used. The statistical significance of these coefficients was assessed via nonparametric bootstrapping with 1,000 replicates. The coefficient is deemed significant if the 95% confidence interval does not include zero.

#### *1.11 Unsupervised clustering analysis*

Given that it is usually difficult to obtain robust estimates of population census size ($N_c$) and life history traits, we also performed an unsupervised hierarchical clustering analysis on all species to evaluate how well threatened and non-threatened species can be separated using only genetic features without any other ecological features and census size. Firstly, we used normalized continuous genetic features (historical effective population size estimated by SMCPP, genome size, GC content and genetic diversity) as input. Considering the influence of phylogenetic relationships, the standardized genetic data were processed using the 'phyl.pca' function (with "method = 'lambda' ") in the R package 'phytools' (v. 1.5.2) (*41*). Then, the all-PC portions were used for unsupervised clustering, which to some extent eliminated the influence of phylogenetic signals. Finally, the optimal number of clusters was determined using the 'fviz_nbclust' function in the R package 'factoextra' (v. 1.0.7) (*42*).

### 2. Results

#### *2.1 Genomic datasets*

For 101 species in this study, 83 have reference genomes available either from the public domain or assembled in this study. For the remaining 18 species, reference genomes of 14 closely related species from the same genera of the focus species were used for short read alignment and the genomic architecture statistic calculation. These included *Begonia masoniana* (for *B. picturata*, and *B. rhynchocarpa*), *Camellia chekiangoleosa* (for *C. impressinervis*, *C. pingguoensis* var. terminalis, and *C. pubipetala*), *Capsella rubella* (for *C. grandiflora*), *Coptis chinensis* (for *C. quinquesecta*), *Dioscorea zingiberensis* (for *D. nitens*), *Fagus sylvatica* (for *F. hayatae* and *F. orientalis*), *Helianthus annuus* (for *H. giganteus*), *Hopea hainanensis* (for *H. reticulata*), *Magnolia sinica* (for *M. odoratissima*), *Pinus tabuliformis* (for *P. wangii*), *Plantago major* (for *P. lanceolata*), *Quercus dentata* (for *Q. aliena*), *Salweenia bouffordiana* (*S. wardii*), and *Zea mays* (for *Z. diploperennis*). The reference genomes exhibited high assembly completeness with a mean BUSCO score equal to 96.92% (including both single-copy and duplicated genes) in genome mode. The genomic architectures, including genome size (mean = 1.69 Gb), GC content (mean = 35.69%), repetitive sequence content (mean = 53.81%), and gene count (mean = 37,067) are summarized across all reference genomes in **Table S5** and the distributions were illustrated in **Fig. S10**. In particular, the six *de novo* assembled genomes had an average of BUSCO scores of completeness equal to 98.06% for the genome assembly and 79.71% for protein coding gene annotations (**Fig. S11 & S12**). See also in section **2.6** for the effects of reference genomes on all related analyses.

For the population whole-genome resequencing (WGS) data generated in this study or downloaded from public domains, the average sample size was equal to 19 individuals per species. Following a rigorous filtering workflow from short read mapping to genetic variant identifying (**Fig. S8**), a high-quality genomic dataset was generated for all 101 species, which includes in total 10,546 (in *Helianthus giganteus*) to 78,728,309 SNPs (in *Cupressus gigantea*) (**Table S5**).

#### *2.2 Threatened species have lower genetic diversity but higher selection efficacy*

We calculated individual heterozygosity (mean = 0.00523) and sequencing depth (mean = 23.24×) across the dataset (**Fig. S13A & B**, **Table S7** & **S8**). A significant correlation was identified between heterozygosity and sequencing depth ($P$ < 0.001), however, the effect is negligible with adjusted $R^2$ equal to 0.03 (**Fig. S12C**). Subsequently, we computed five key diversity metrics for each species: 1) genome-wide overall nucleotide diversity ($\pi$); 2) Watterson's $\theta$; 3) species-level heterozygosity; 4) nucleotide diversity at zero-fold ($\pi_0$) and 5) nucleotide diversity at four-fold ($\pi_4$) degenerate sites. These metrics exhibited substantial variation, spanning two to three orders of magnitude (**Table S1**). The ratio of non-synonymous to synonymous nucleotide diversity ($\pi_0/\pi_4$) was derived as a proxy for the efficacy of selection. Given the highly positive correlations observed among all five genetic diversity metrics (**Fig. S6**), we retained neutral nucleotide diversity ($\pi_4$) as the representative metric for all subsequent analyses to avoid redundancy. In general, threaten species have significantly lower $\pi_4$ but higher $\pi_0/\pi_4$ ratio (see **Fig. 2** in main text).

#### *2.3 Threatened species have smaller $N_e$, but larger $N_e$ changes and genome sizes*

Previous studies have demonstrated that threatened species generally exhibit specific genetic features, including reduced effective population sizes ($N_e$), low genetic diversity, and elevated $N_e/N_c$ ratio (*43-45*). A recent study further suggested a positive association between extinction risk and genome size (*46*). Our analysis strongly corroborates these hypotheses: compared to non-threatened and Data Deficient (DD) categories, threatened species in our dataset displayed significantly lower $\widehat{N_e}$ (across both ancient and recent timescales) and neutral nucleotide diversity. Conversely, they exhibited significantly higher genome sizes and $\widehat{N_e}/\widehat{N_c}$ ratios, a pattern consistent across temporal stages (**Fig. S3**).

Threatened species consistently exhibited reduced $\widehat{N_e}$ across all temporal scales. Ancient $\widehat{N_e}$ ($10^5$–$10^7$ generations ago) was significantly lower in threatened species (median = 345,000) compared to non-threatened (median =1,190,000, p-value = 0.01) and DD groups (median = 800,000) (**Fig. S3**). This disparity continued into recent period (1–200 generations ago), where $\widehat{N_e}$ for threatened species (median = 1652) fell significantly below both non-threatened (median = 3611, p-value = 0.04) and DD species (median = 3462) (**Fig. S3).**

Conversely, $N_e/N_c$ ratios displayed an inverse pattern. Across both two estimators (EPOS and GONE2), values were significantly higher in threatened species (medians: EPOS = 235.50; GONE2 = 0.61) relative to non-threatened (EPOS = 1.18, p-value = $1.59\times10^{-6}$; GONE2 = 0.0017, p-value = $3.91\times10^{-9}$) (**Fig. S3**). This elevated ratio in threatened species was also significant when compared to DD group (EPOS = 3.58, p-value = $1.09\times10^{-3}$; GONE2 = 0.02, p-value = $1.08\times10^{-4}$).

Meanwhile, threatened species experienced a more severe decline in historical $N_e$ estimated by SMCPP (median = 97.9%, range: 81.2%-99.7%) compared to both non-threatened (median = 93.6%, range: 6.8%-99.8%, not significant) and DD species (median = 90.5%, range: 71.4%-99.9%, p-value = 0.04) (**Fig. S3**). Finally, conforming to theoretical expectations, threatened species had reduced genetic diversity (median = $4.15\times10^{-3}$) but larger genome sizes (median = 0.91 Gb）. These metrics differed significantly from those of non-threatened ($\pi_4$ median = $7.60\times10^{-3}$, p-value = $3.88\times10^{-4}$; genome size median = 0.72 Gb) and DD ($\pi_4$ median = $9.31\times10^{-3}$, p-value = $1.50\times10^{-4}$; genome size median = 0.41 Gb, p-value = $2.22\times10^{-5}$) (**Fig. S3**).

#### *2.4 Genetic features are good predictors of conservation status*

First, based on Random Forest results (**Fig. 4D** in main text), we classified three Data Deficient (DD) species *Populus qiongdaoensis*, *Begonia rhynchocarpa*, and *B. picturata* as 'threatened', and the remaining 24 DD species as 'non-threatened'. The classification of DD species have also been confirmed by the extinction risk predictions carried out by Bachman et al., (2024) based on approximately 330,000 angiosperm species (*47*). This classification yielded a final dataset of 52 threatened and 49 non-threatened species for downstream clustering and demographic regressions.

Unsupervised clustering (*K* = 2) effectively separated the taxa into a 'genetically threatened' clade (n = 52; red branches in main text **Fig. 6**) and a 'genetically non-threatened' clade (n = 49; green branches). The overall clustering accuracy of the model is equal to 69.31%, which indicates that genetic features can be good predictors of conservation status even if census population size and other ecological features are not available. However, the model had a high Type I error rate (false positives) equal to 61.22%, but a rather low Type II error rate (false negatives) equal to 1.92%.

#### *2.5 The effects of census size and genetic diversity on conservation classification*

Model 4 was selected as the best-fitting path model with the difference of ΔCICc larger than 7.8, and a dominant weight ($\omega$ = 0.968) over all others (**Table S9**). All effects in Model 4 also passed the directed separation tests ($P$ = 0.561; **Fig. S14, Table S10**). Coefficients for paths absent in a given model were assumed to be zero. We extracted the standardized path coefficients and 95% confidence intervals (CIs) from the best model to evaluate the direction and significance of direct effects (**Table S11**).

In the best-fitting model, estimated $\widehat{N_c}$ and $\pi_4$ both exerted significant negative direct effects on conservation status ($\beta$ = -0.42 and $\beta$ = -0.32, respectively) (**Fig. S15**, **Table S11**). Regarding predictors of genetic diversity, SMCPP estimated historical $\widehat{N_e}$, GC content, mating system and $\widehat{N_c}$ all demonstrated significant positive effects on $\pi_4$ ($\beta$ = 0.53, 0.36, 0.31 and 0.22, respectively). Additionally, historical $\widehat{N_e}$ positively affected contemporary $\widehat{N_c}$ ($\beta$ = 0.28). We also identified a significant negative effect of mating system on GC content ($\beta$ = -0.15) and $\widehat{N_c}$ ($\beta$ = -0.16) (**Fig. S15**, **Table S11**).

Overall, the path analysis identified four distinct pathways influencing $\pi_4$ and two primary pathways determining conservation status. Notably, $\widehat{N_c}$ exhibited a larger effect size on conservation status than $\pi_4$, suggesting that while census size is the most direct predictor, estimates of genetic diversity provide a crucial alternative for assessing extinction risk when census data are unavailable.

#### *2.6 Effects of reference genomes*

As specified in **1.6** for 18 species reference genomes of closely related species in the same genera were used to calculate genomic architecture related features (i.e., GC content, genome size, and repetitive content), we re-ran all analyses with these three features for other 83 species for which reference genomes are available. In general, main results were retained (**Fig. S16**). The effect of mating system on genetic diversity became less significant with 95% CI overlapped with zero (**Fig. S16B**). For the 'randomForest' model, *Populus qiongdaoensis* was still predicted to be of high risks of extinction. Two Begonia species (*B. rhynchocarpa* and *B. picturata*) were no longer in this list due to lacking of reference genome (**Fig. S16F**). Population census size and genetic diversity were also the main predictors of conservation status in the phylogenetic path analysis (**Table S12**). Finally, for the unsupervised clustering analysis based on genetic and ecological features, the results were still highly correlated to IUCN conservation status (Fisher's exact test p-value = $3.127\times10^{-4}$). Using GC content calculated from reference genomes of focal species can improve model performance by both increasing the accuracy (from 69.31% to 73.49%) and reducing Type I error (from 61.22% to 40.91%). On the other hand, Type II error was increased from 1.92% to 10.26%, possibly due to fewer sample size (**Fig. S16H**), which means GC content may be conserved across some closely related species and can provide some information for prediction.

To summarize, using reference genome from a different species can lead to reduced prediction accuracy mainly by increasing false positive rate. If the genomic features are conserved between focal and related species (like GC content), it may also reduce the false negative rate and lead to the conserved results for potential threatened species.

**Supporting figures**

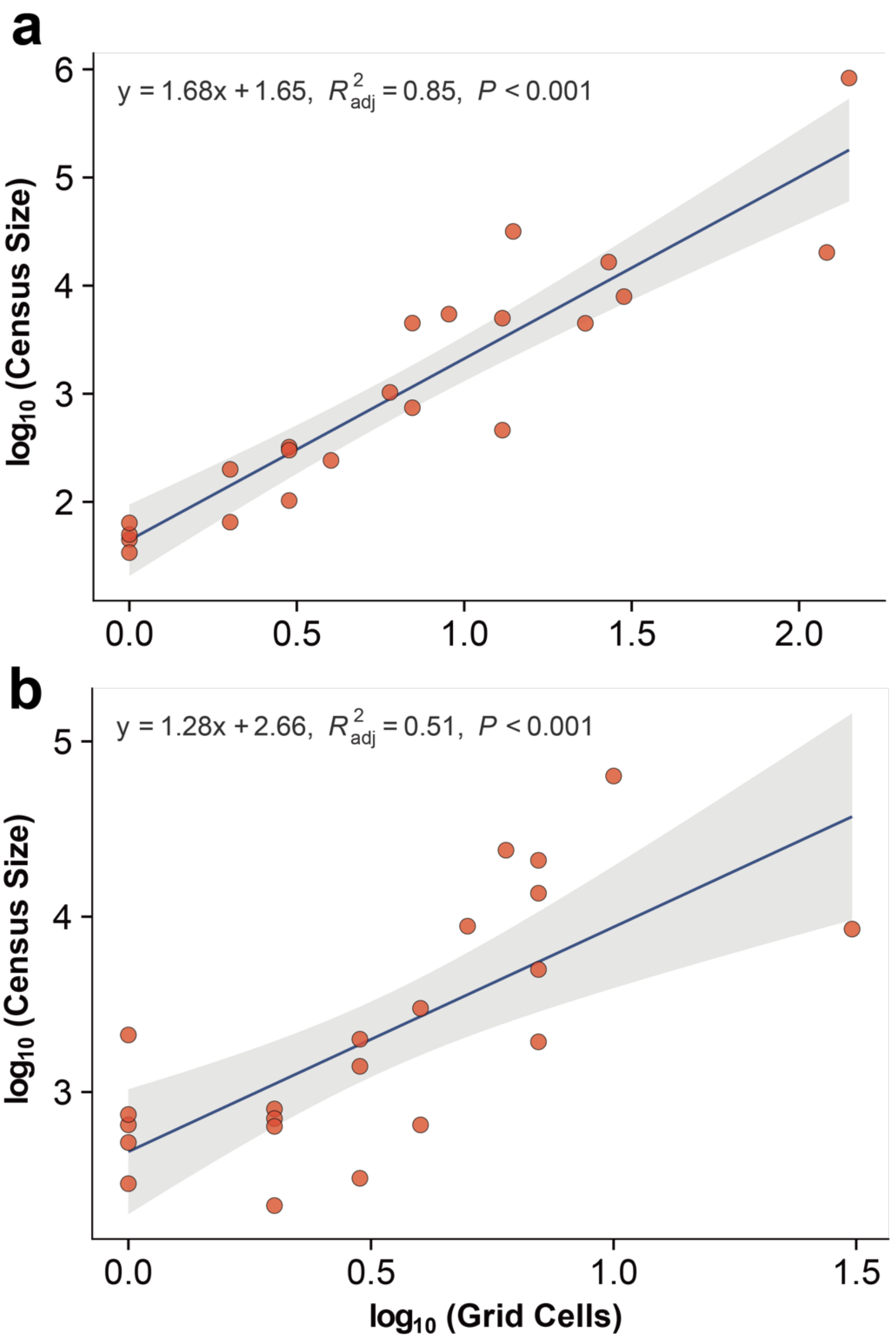


**Fig. S1. Comparison of census population size estimation methods for (a) tree, (b) shrubs and herbaceous species.** Solid lines represent the linear fit, shaded gray areas indicate 95% confidence intervals, and red dots denote individual species.

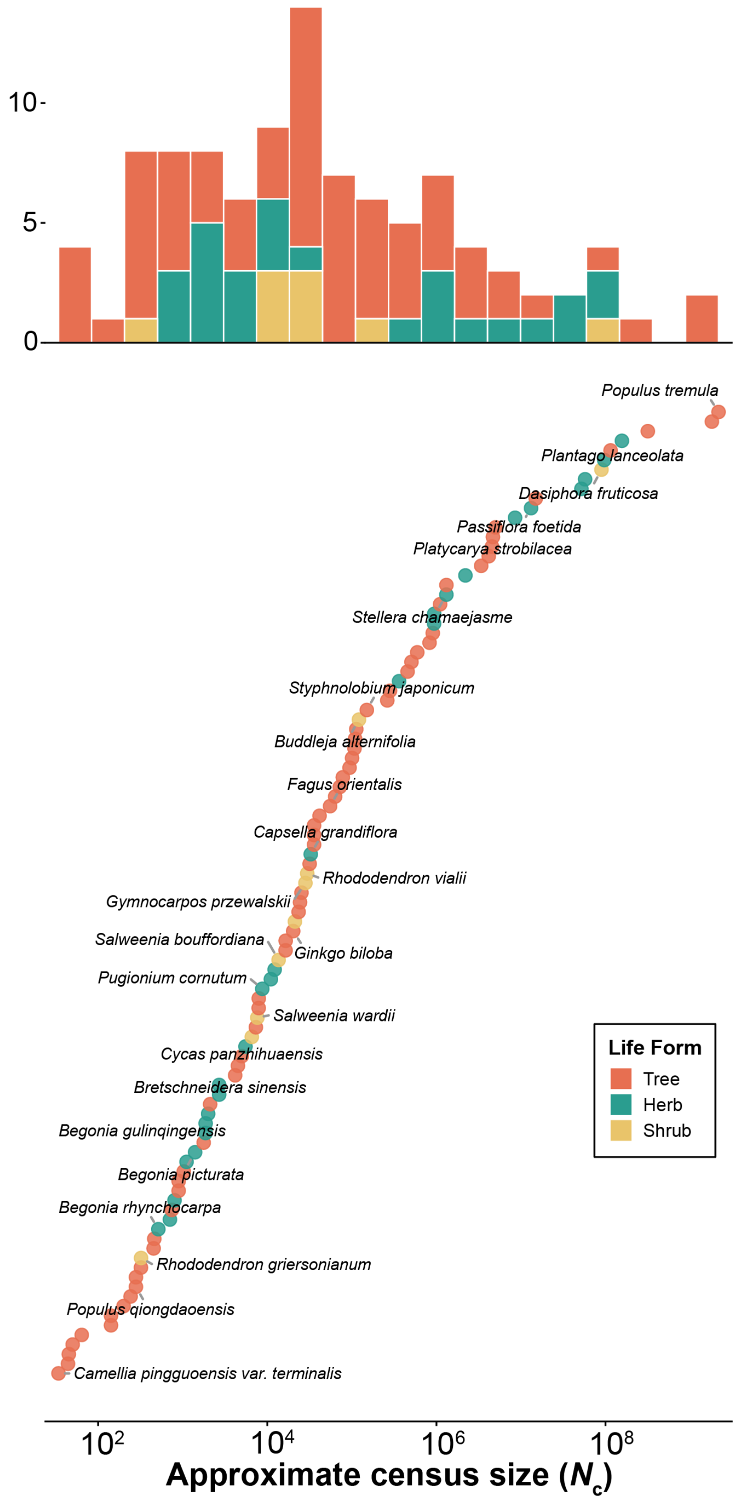


**Fig. S2.** The distribution of approximate census population sizes estimated by this study.

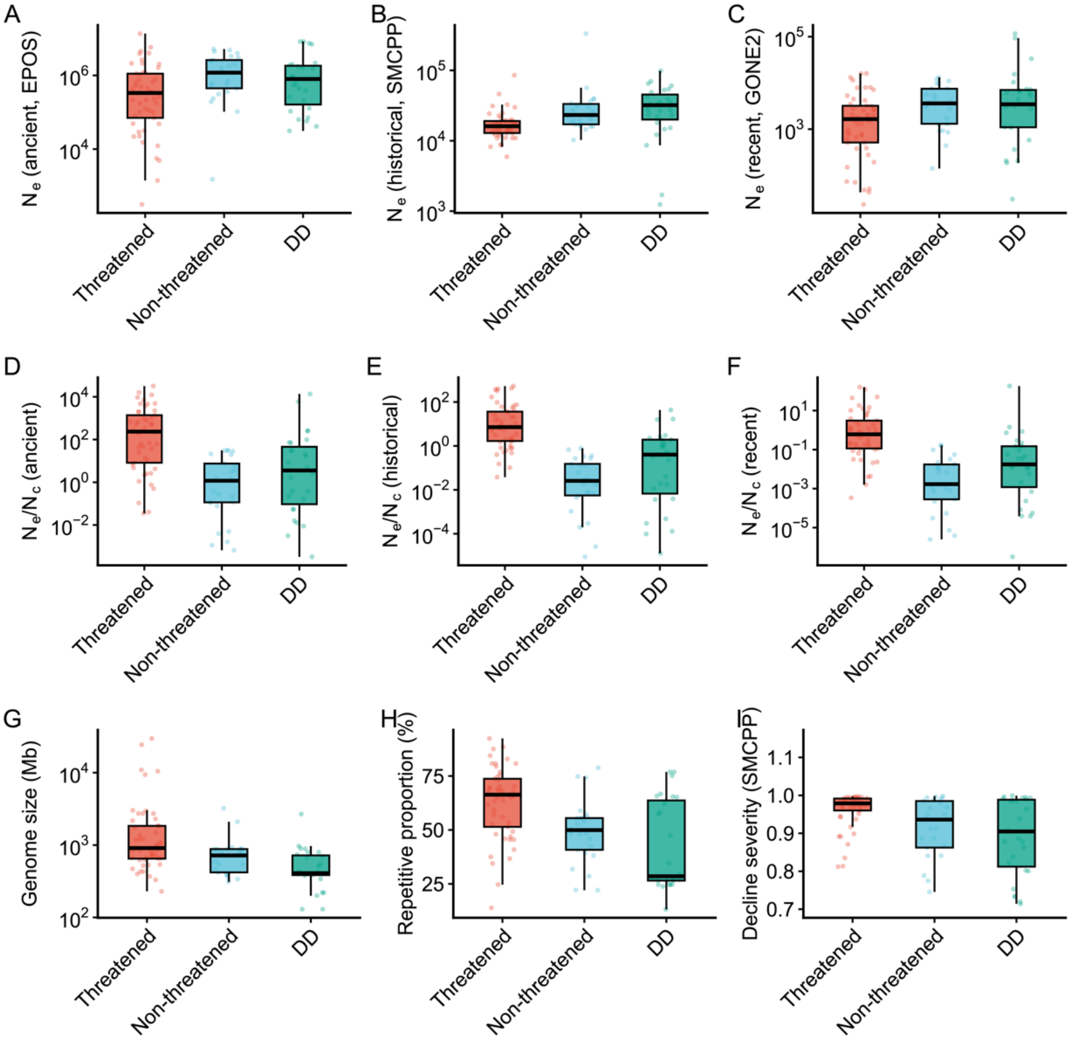


**Fig. S3. The distributions of demographic and genomic metrics across conservation categories.** (**A-C**) The distributions of ancient, historical, and recent effective population size based on EPOS, SMCPP, and GONE2, respectively; (**D-F**) The distributions of $N_e/N_c$ ratio with corresponding $N_e$ estimated by EPOS, SMCPP, and GONE2, respectively; (**G**) the distributions of genome size; (**H**) the percentage of repetitive elements; (**I**) the distributions of SMCPP-based $N_e$ decline severity.

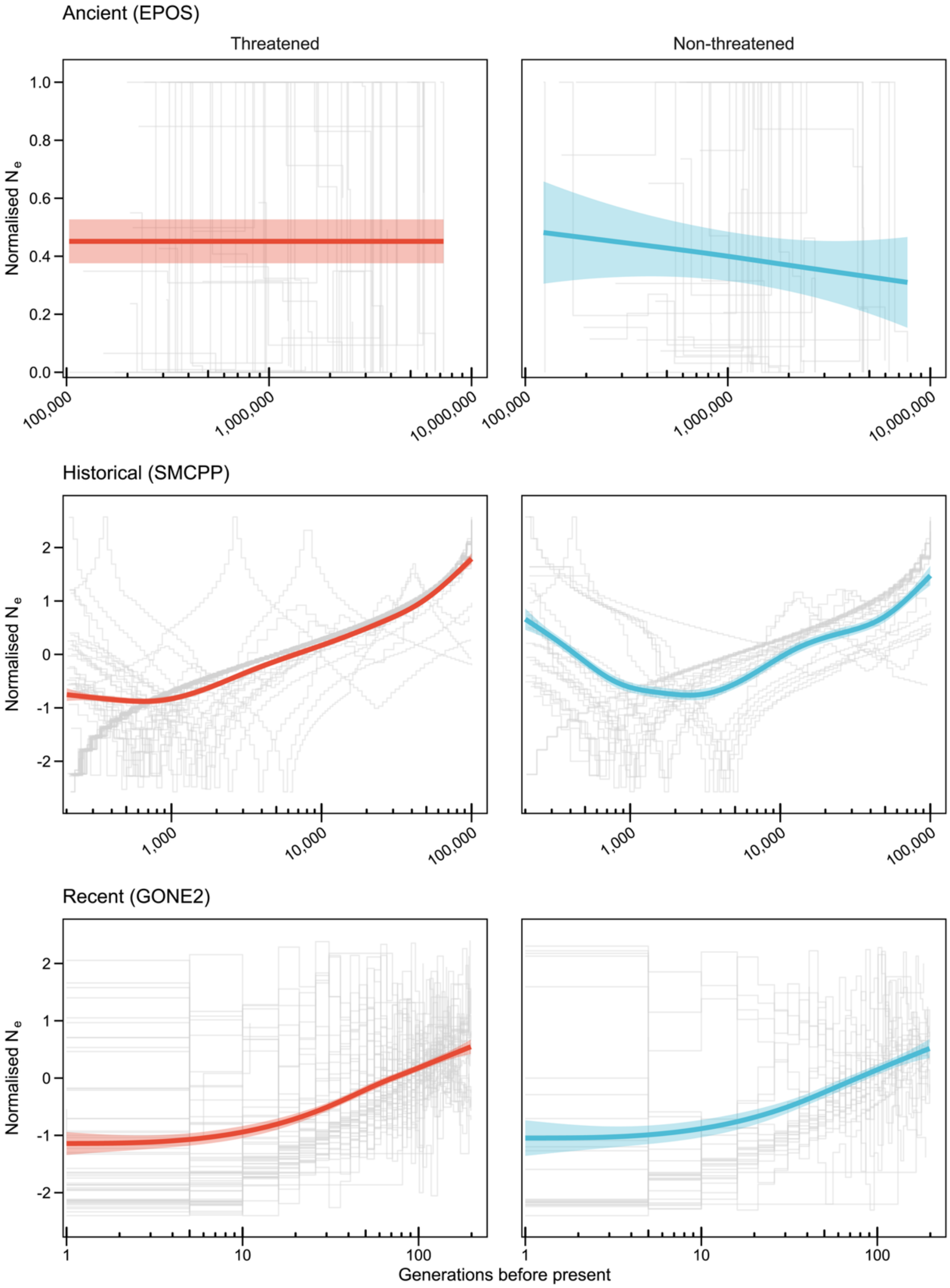


**Fig. S4. Regression analysis of $N_e$ fluctuations over time in generations**. Standardized $N_e$ trajectories for 52 threatened and 25 non-threatened species across ancient (EPOS), historical (SMCPP), and recent (GONE2) time periods (top to bottom). Gray lines indicate individual species trajectories. Red and blue lines represent the generalized additive model (GAM) fitted curves for threatened and non-threatened groups, respectively. See section **1.10** for standardization details.

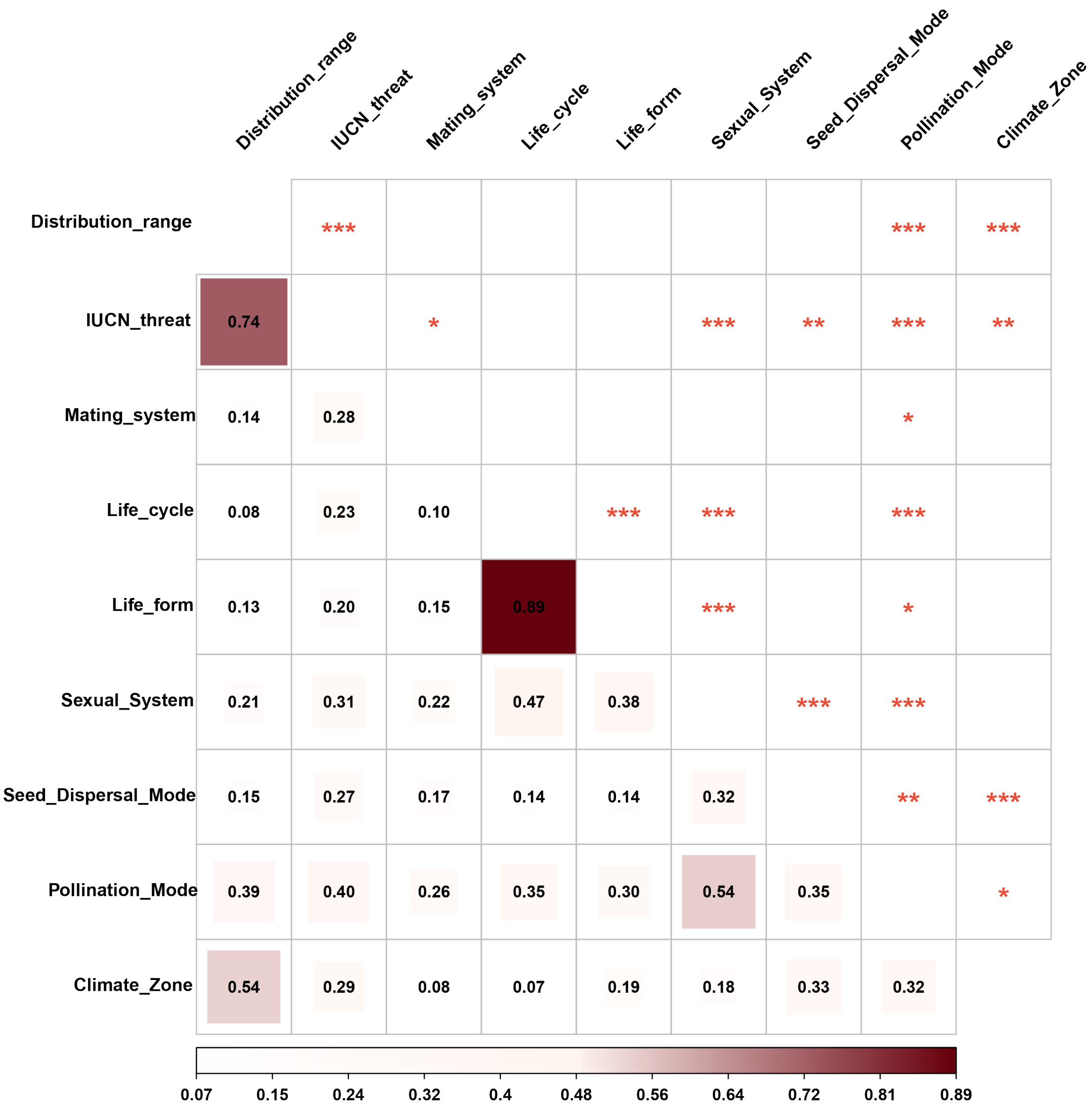


**Fig. S5. Pairwise correlation heatmap of categorical variables**. Analysis performed across 101 species. Distribution range contains ‘wide’ and ‘narrow’ two types. The lower triangle displays the correlation coefficients, represented by both numerical values and box sizes. The upper triangle indicates statistical significance. The diagonal is masked. Significance levels: **P* < 0.05; ***P* < 0.01; ****P* < 0.001.

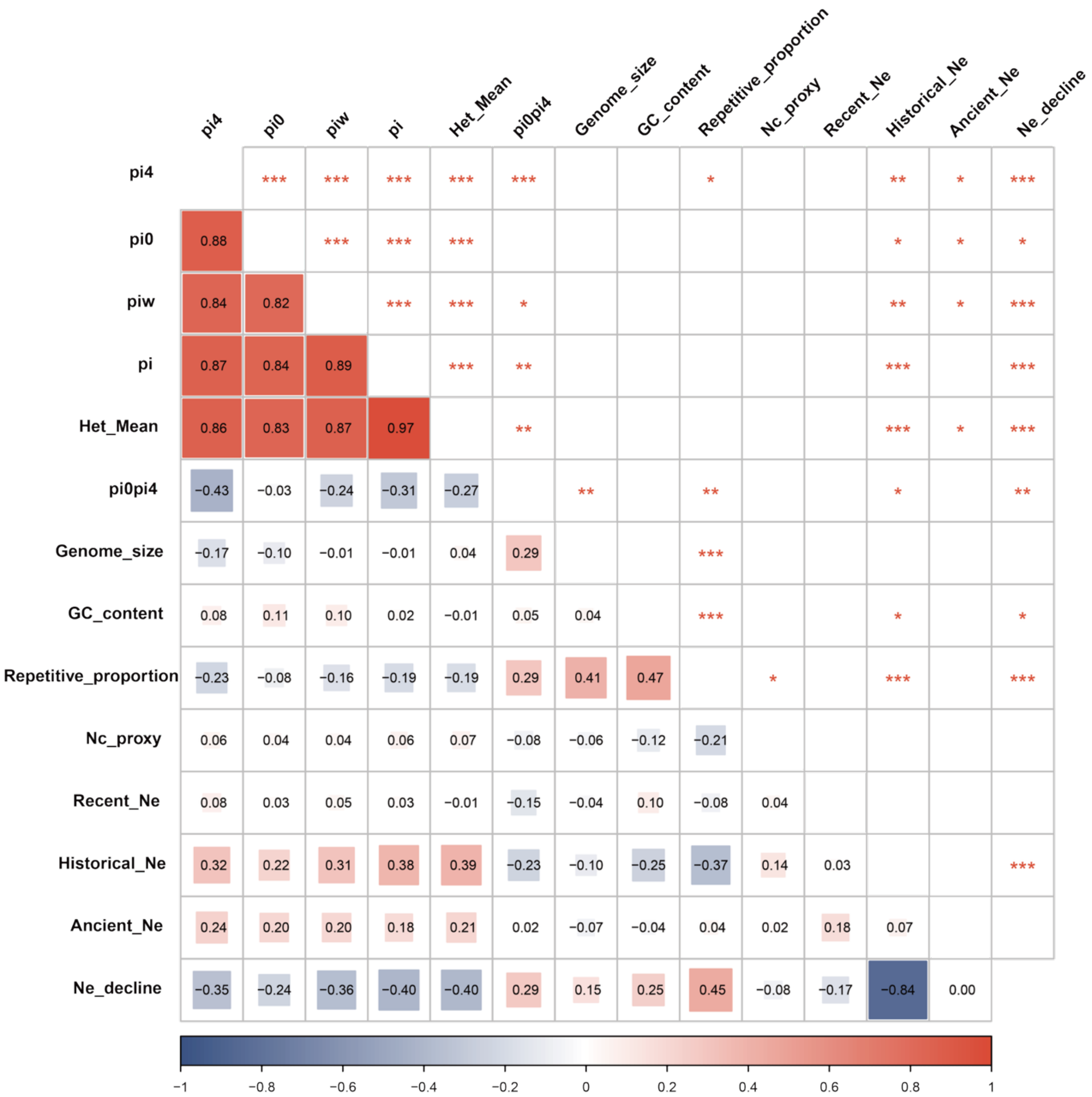

**Fig. S6. Pairwise correlation heatmap of continuous variables**. Analysis performed across 101 species. The lower triangle displays the correlation coefficients, represented by both numerical values and box sizes. The upper triangle indicates statistical significance. The diagonal is masked. Significance levels: *$P < 0.05$; **$P < 0.01$; ***$P < 0.001$. Variable abbreviations are listed: **pi4**: nucleotide diversity at 4-fold degenerate sites; **pi0**: nucleotide diversity at 0-fold degenerate sites; **piw**: Watterson's $\theta$; **pi**: genome-wide diversity; **Het_Mean**: mean species-level heterozygosity; **pi0pi4**: the ratio of synonymous to non-synonymous nucleotide diversity; **Ancient** $N_e$: harmonic mean $N_e$ ($\widehat{N_e}$) estimated by EPOS in interval [$10^5$-$10^7$ ] generations; **Historical** $N_e$: $\widehat{N_e}$ estimated by SMCPP in interval [200-100,000 generations]; $N_e$ **decline**: decline severity of $N_e$ estimated by SMCPP; **Recent** $N_e$: $\widehat{N_e}$ estimated by GONE2 in interval [1, 200] generations.

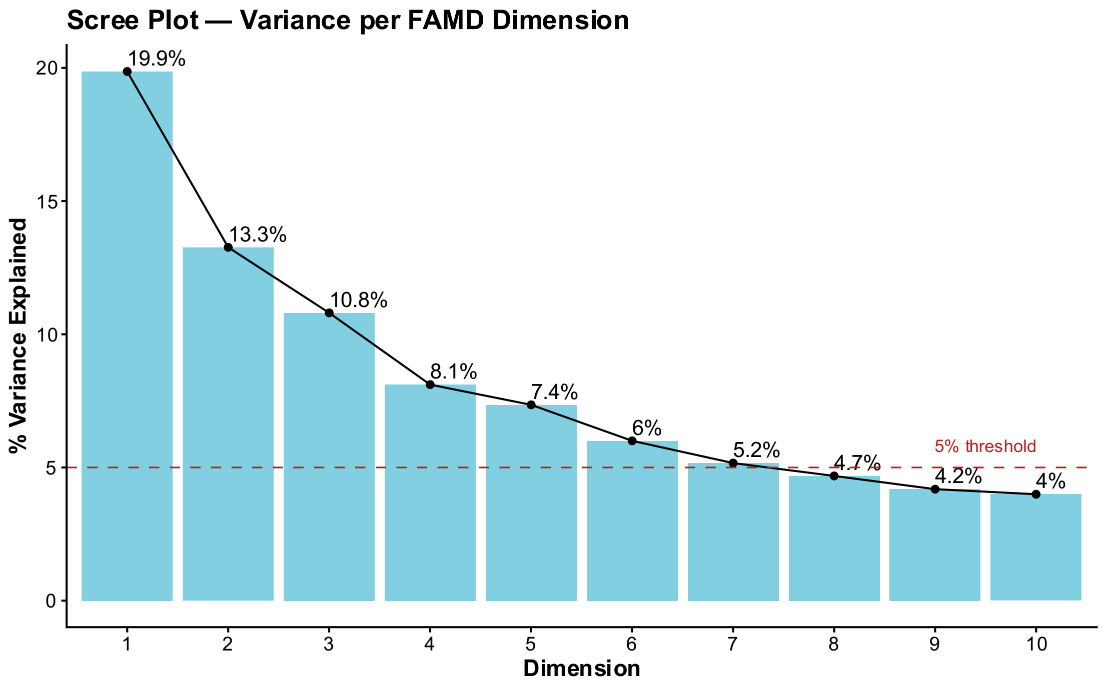


**Fig. S7. Scree plot of variance explained**. The bar plot displays the proportion of total variance captured by each of the first ten principal components (PCs) derived from the FAMD analysis of 101 species.

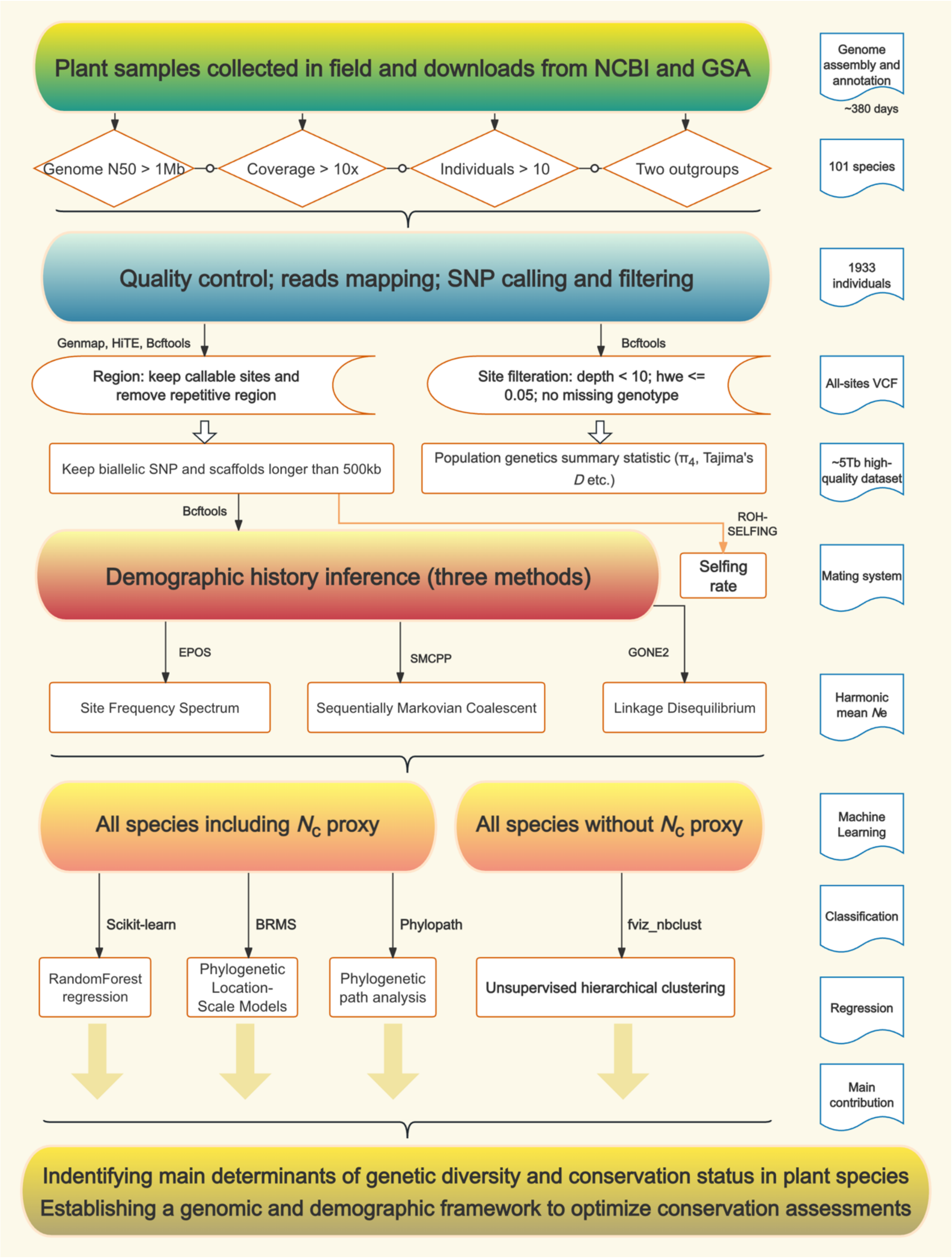


**Fig. S8**. **The workflow used in the study**.

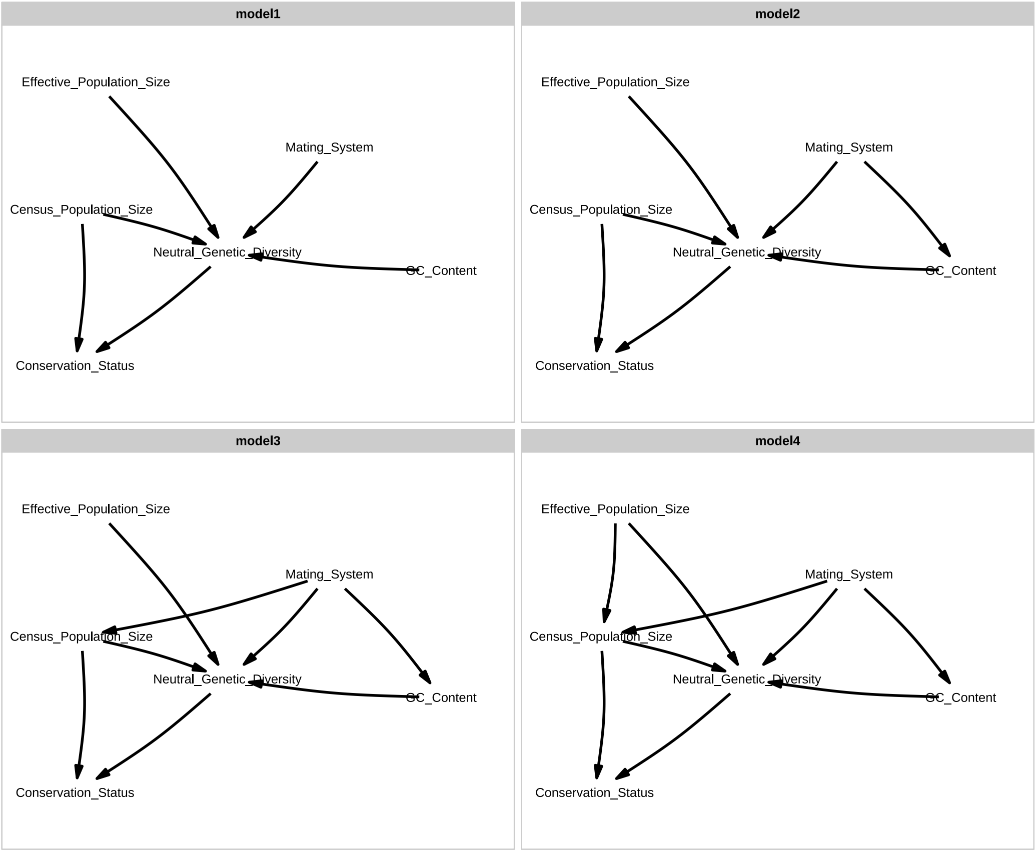


**Fig. S9. Four candidate structural models were evaluated via phylogenetic path analysis**.

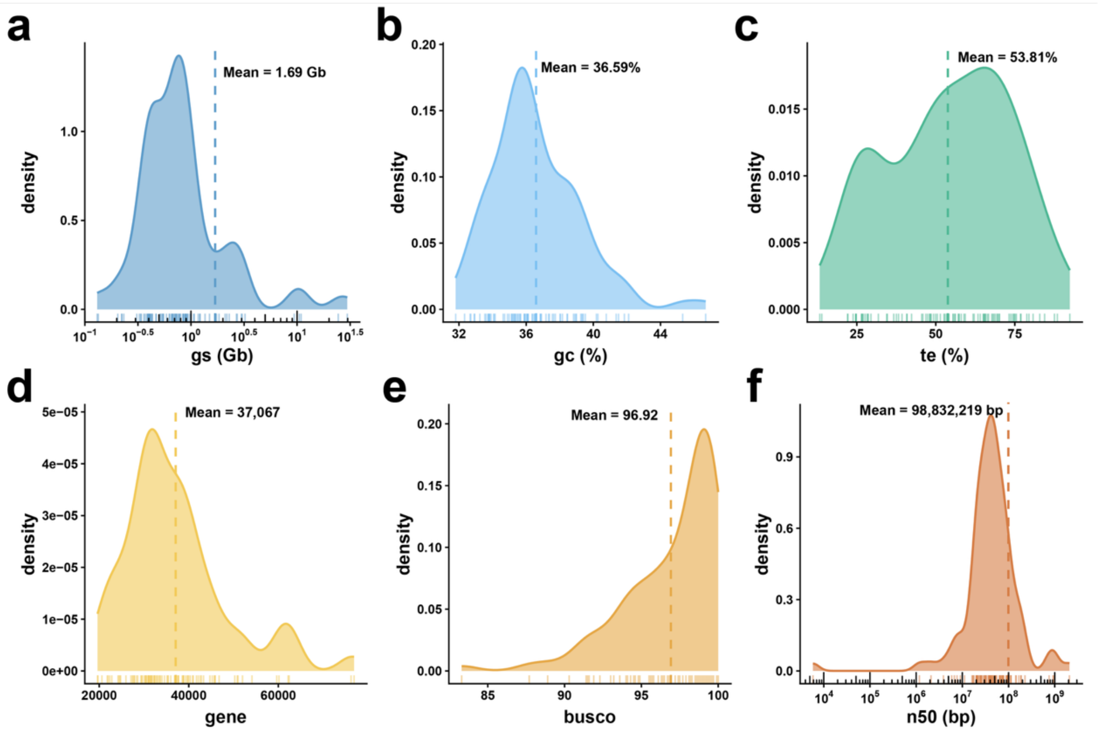


**Fig. S10**. **Schematic diagram of genomic statistical metrics of 101 plant species in the study**. a) Distribution of genome size; b) Distribution of GC content; c) Distribution of repetitive proportion within genomes; d) Distribution of gene numbers across species genomes; e) Distribution of BUSCO completeness scores; f) Distribution of N50 values across genomes. The dotted line indicates the average value.

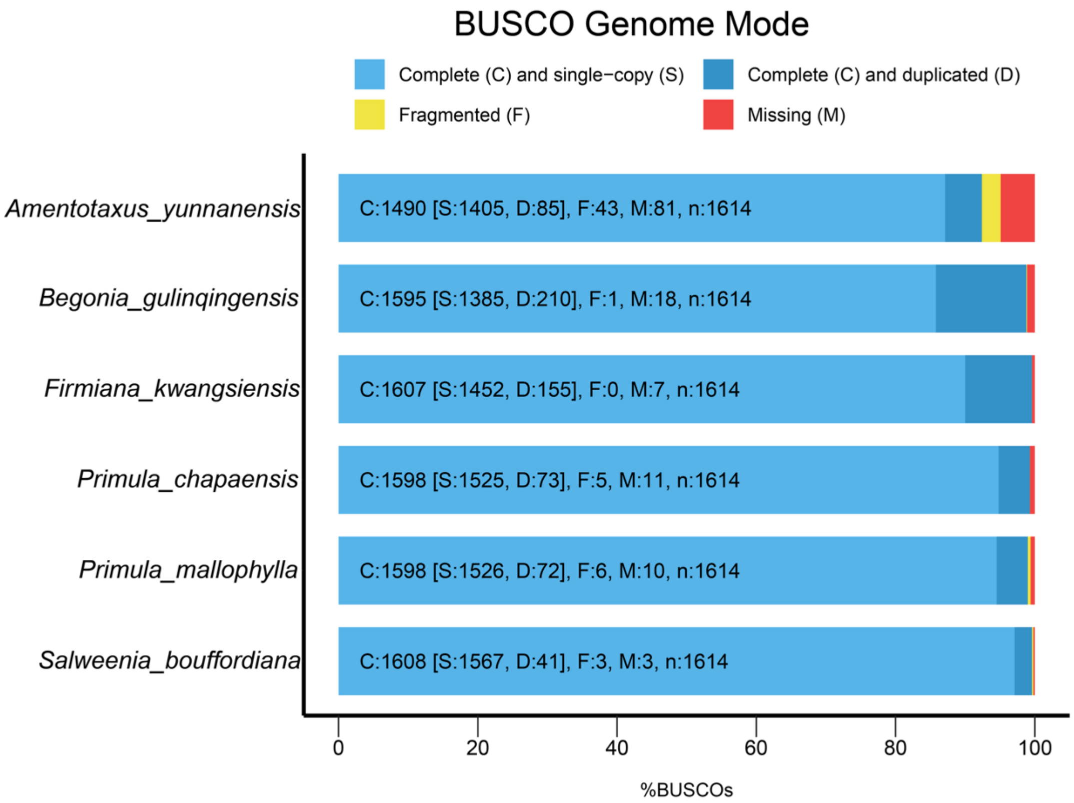


**Fig. S11**. **BUSCO of assembly against the Embryophyta_odb10 database**.

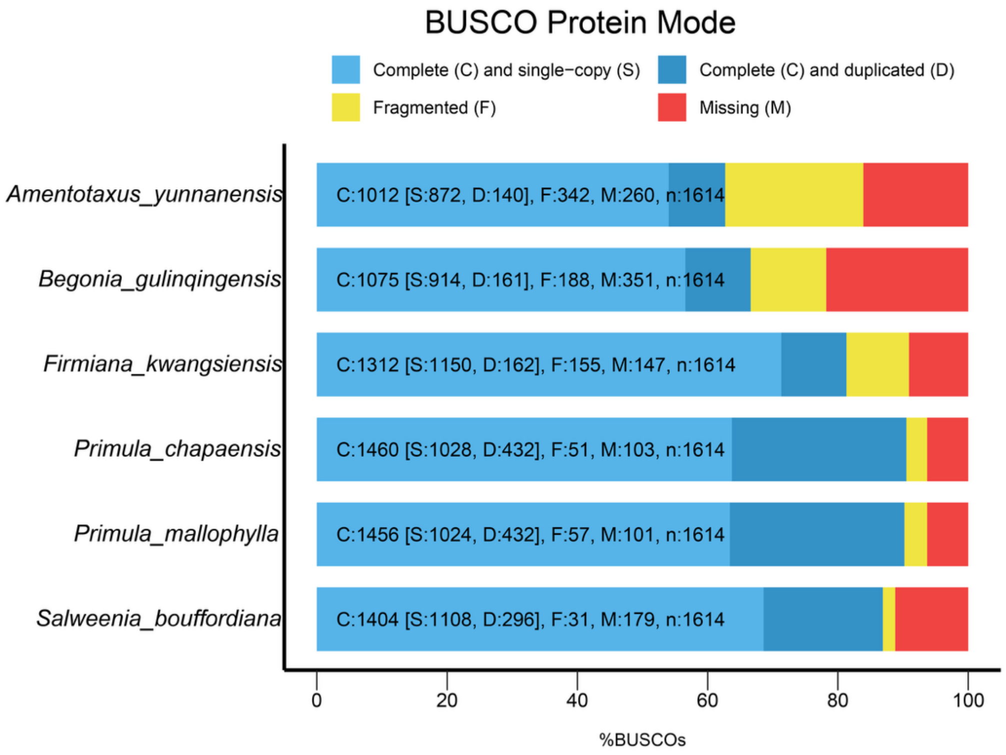


**Fig. S12**. **BUSCO of protein-coding genes against the Embryophyta_odb10 database**.

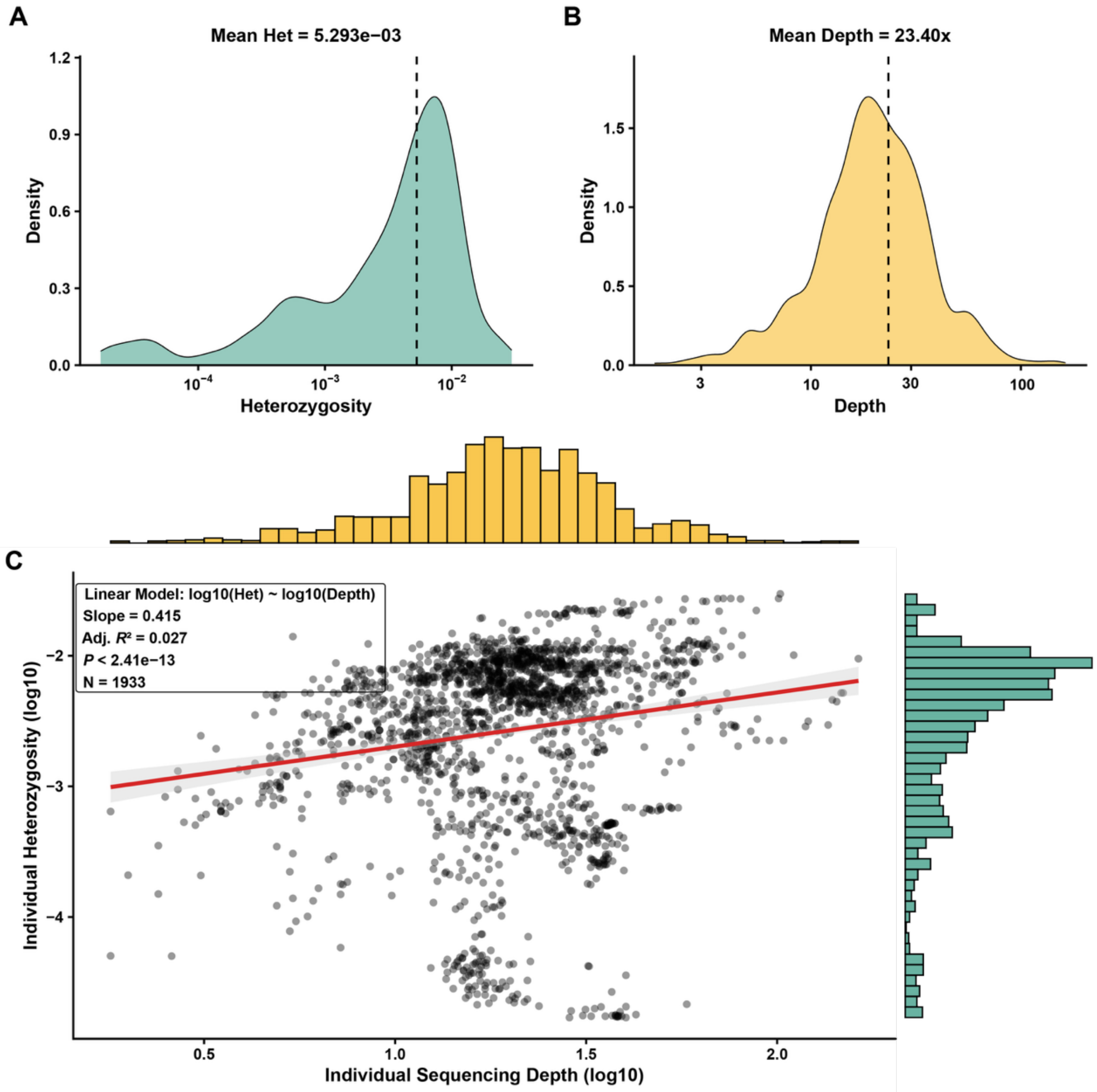


**Fig. S13**. **The distribution and correlation between heterozygosity and depth in individual-level**. (**A**) the distribution of individual heterozygosity. (**B**) the distribution of individual sequencing depth. The dotted line indicates the average value. (**C**) the correlation between heterozygosity and depth. Histograms of the periphery show trends in the distribution of data. The solid red line is the line of the linear fit, the grey area indicates the 95% confidence interval, and each species is indicated by a grey dot.

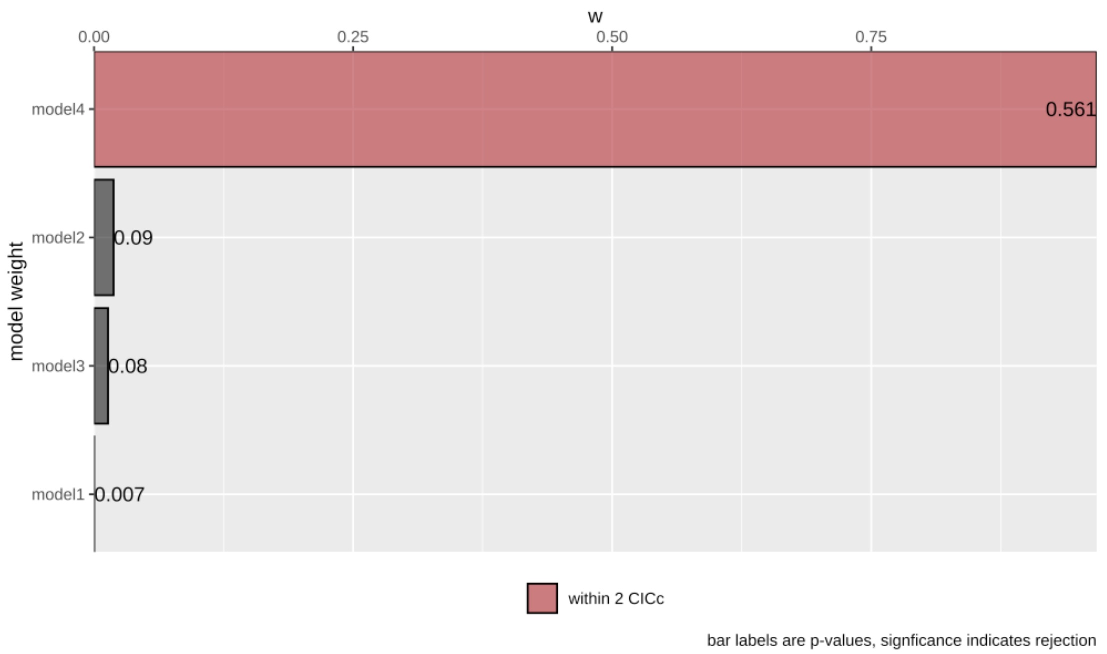


**Fig. S14. Model selection for the phylogenetic path analysis**. Bar lengths represent the relative model weights (**w**) for the four candidate structural models, calculated based on the C-statistic Information Criterion (**CICc**). The best performing model (model4) is highlighted in red, indicating it fall within a **ΔCICc** of 2 from the best-fitting model. The numerical values adjacent to each bar denote the ***p*-values** derived from directed separation tests.

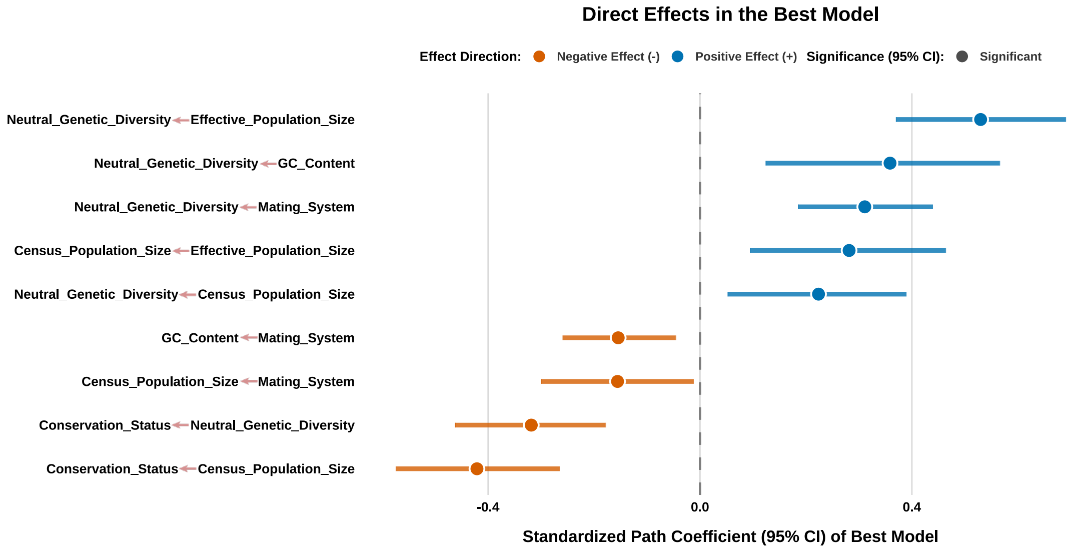


**Fig. S15. Standardized path coefficients for direct effects in the phylogenetic path averaged models**. Points represent the estimated standardized path coefficients for each directed relationship (y-axis), with horizontal lines indicating the 95% confidence intervals (CIs). Point color denotes the direction of the effect: blue indicates a positive effect, while orange indicates a negative effect. Point shape denotes statistical significance: circles indicate significant effects where the 95% CIs does not overlap zero (vertical dashed line), whereas squares indicate non-significant effects where the 95% CIs includes zero.

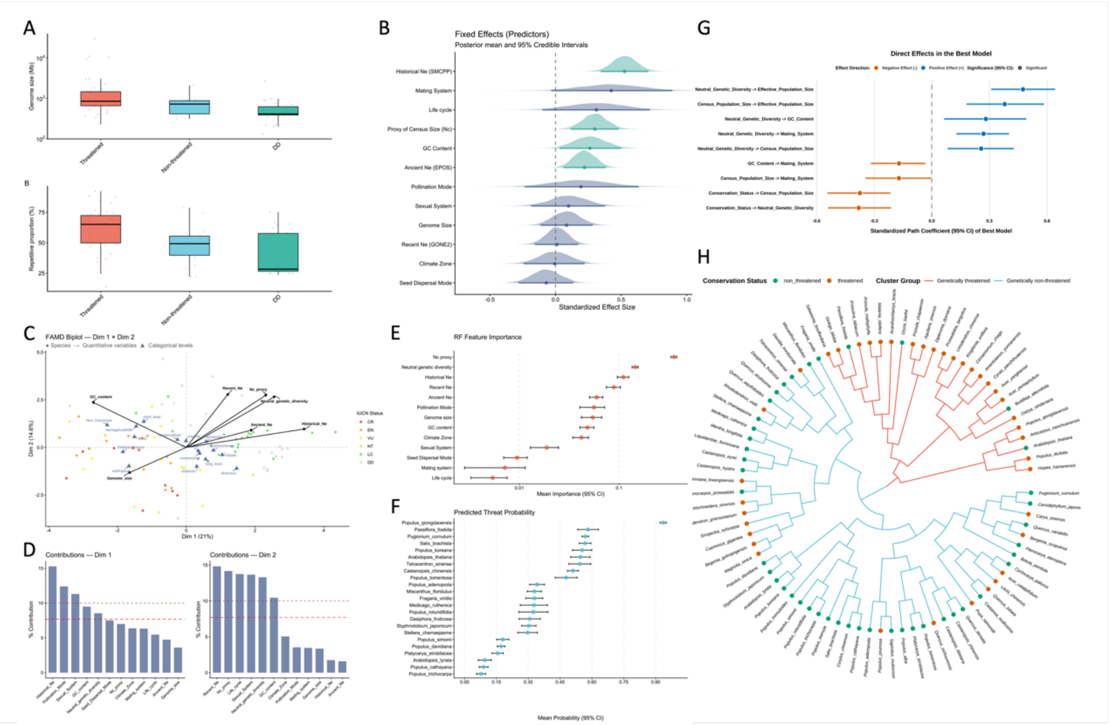

**Fig. S16. Results of analyses re-run in 83 species for which reference genomes are available.** (**A**) The distributions of genome size and repetitive content; (**B**) The distributions of fixed effects of the Bayesian regression model for genetic diversity; (**C**) The FAMD biplot; (**D**) The contribution of each variable to Dim 1 (left) and Dim 2 (right), respectively. (**E**) Feature importance obtained from the 'RandomForest' model; (**F**)The predicted threatened probabilities for Data Deficient (DD) species based on ten random iterations; (**G**) Standardized path coefficients for direct effects in the phylogenetic path averaged models; (**H**) Unsupervised clustering analysis for the prediction of conservation status based on genetic features only

**Supplementary PDF-File “SMCPP-Ne-curve”**

The following figures describe the curves of historical effective population size change for all 101 species based on the SMCPP estimates.

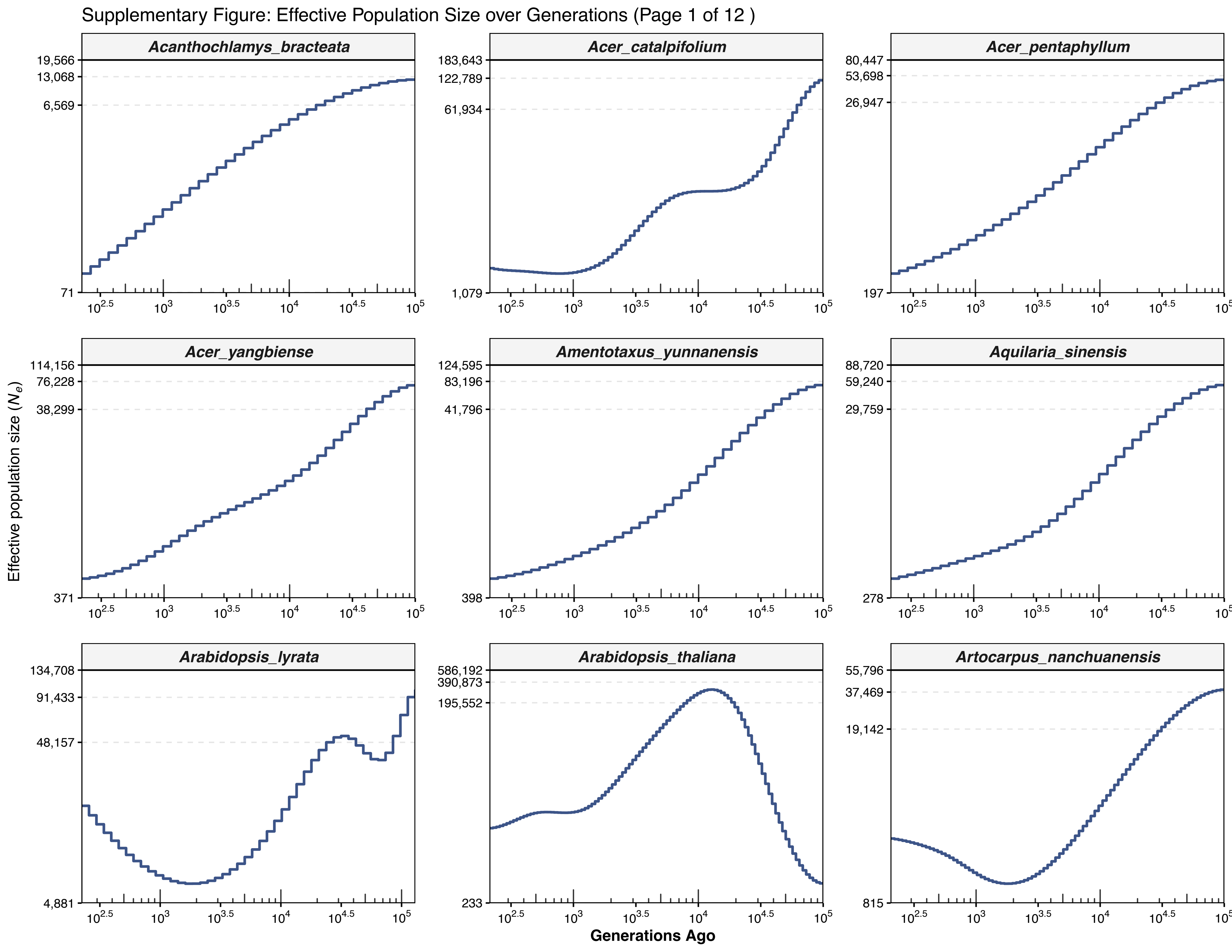
Supplementary Figure: Effective Population Size over Generations (Page 1 of 12 )
Acanthochlamys_bracteata
Acer_catalpifolium
Acer_pentaphyllum
Acer_yangbiense
Amentotaxus_yunnanensis
Aquilaria_sinensis
Arabidopsis_lyrata
Arabidopsis_thaliana
Artocarpus_nanchuanensis
Effective population size ($N_e$)
Generations Ago
19,566
13,068
6,569
71
183,643
122,789
61,934
1,079
80,447
53,698
26,947
197
114,156
76,228
38,299
371
124,595
83,196
41,796
398
88,720
59,240
29,759
278
134,708
91,433
48,157
4,881
586,192
390,873
195,552
233
55,796
37,469
19,142
815
$10^{2.5}$
$10^{3}$
$10^{3.5}$
$10^{4}$
$10^{4.5}$
$10^{5}$

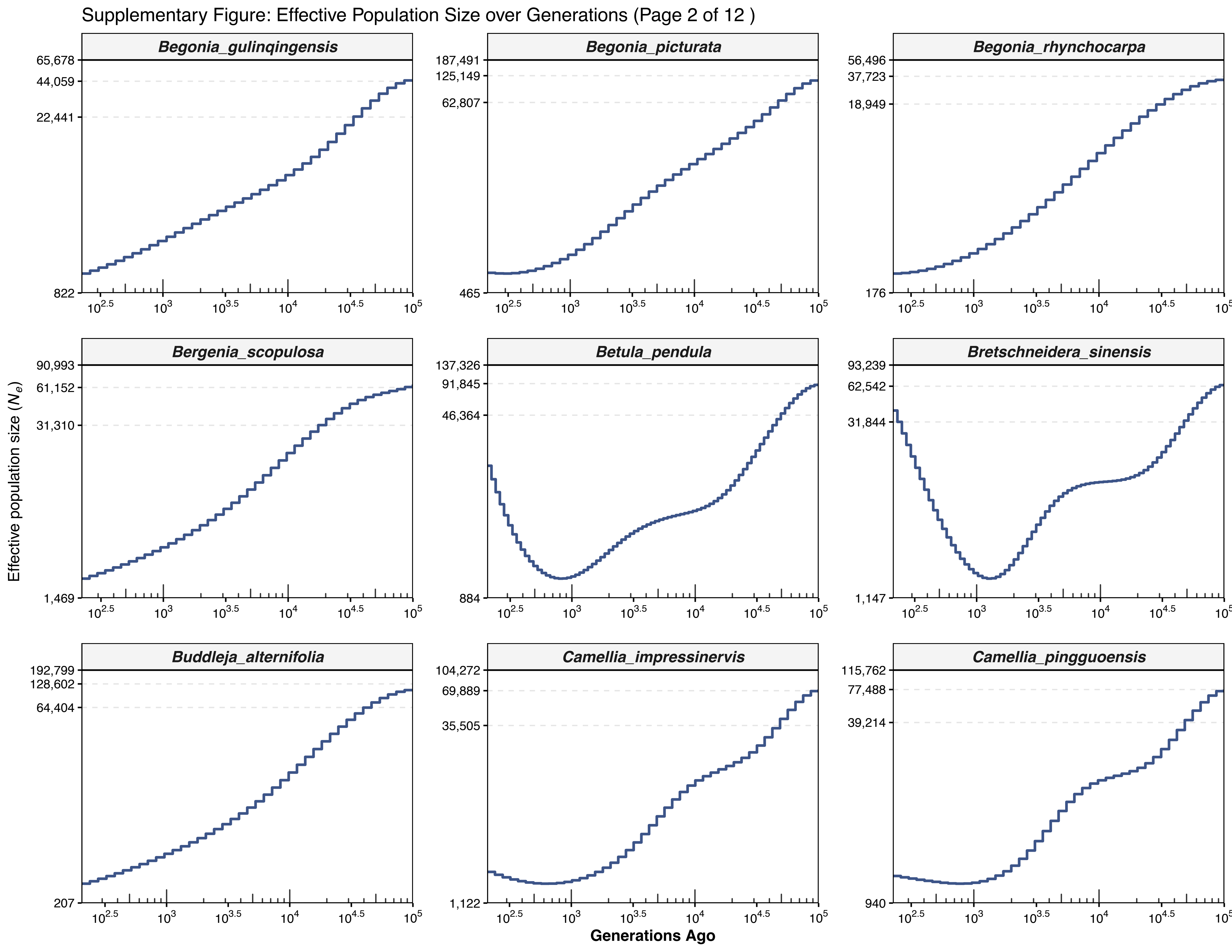

Supplementary Figure: Effective Population Size over Generations (Page 2 of 12 )
Begonia_gulinqingensis
Begonia_picturata
Begonia_rhynchocarpa
Bergenia_scopulosa
Betula_pendula
Bretschneidera_sinensis
Buddleja_alternifolia
Camellia_impressinervis
Camellia_pingguoensis
Effective population size ($N_e$)
Generations Ago

Supplementary Figure: Effective Population Size over Generations (Page 3 of 12 )

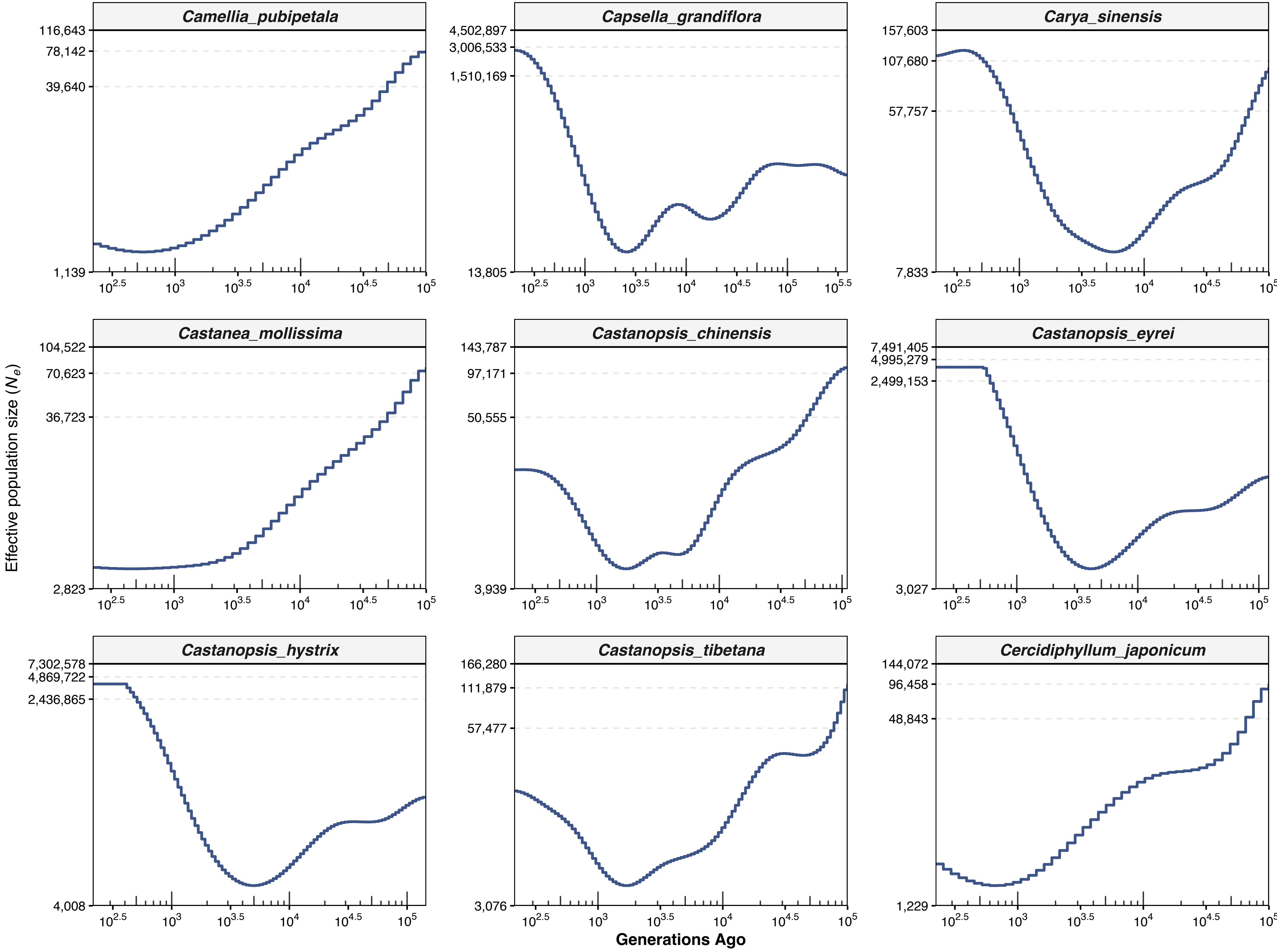

Supplementary Figure: Effective Population Size over Generations (Page 4 of 12 )

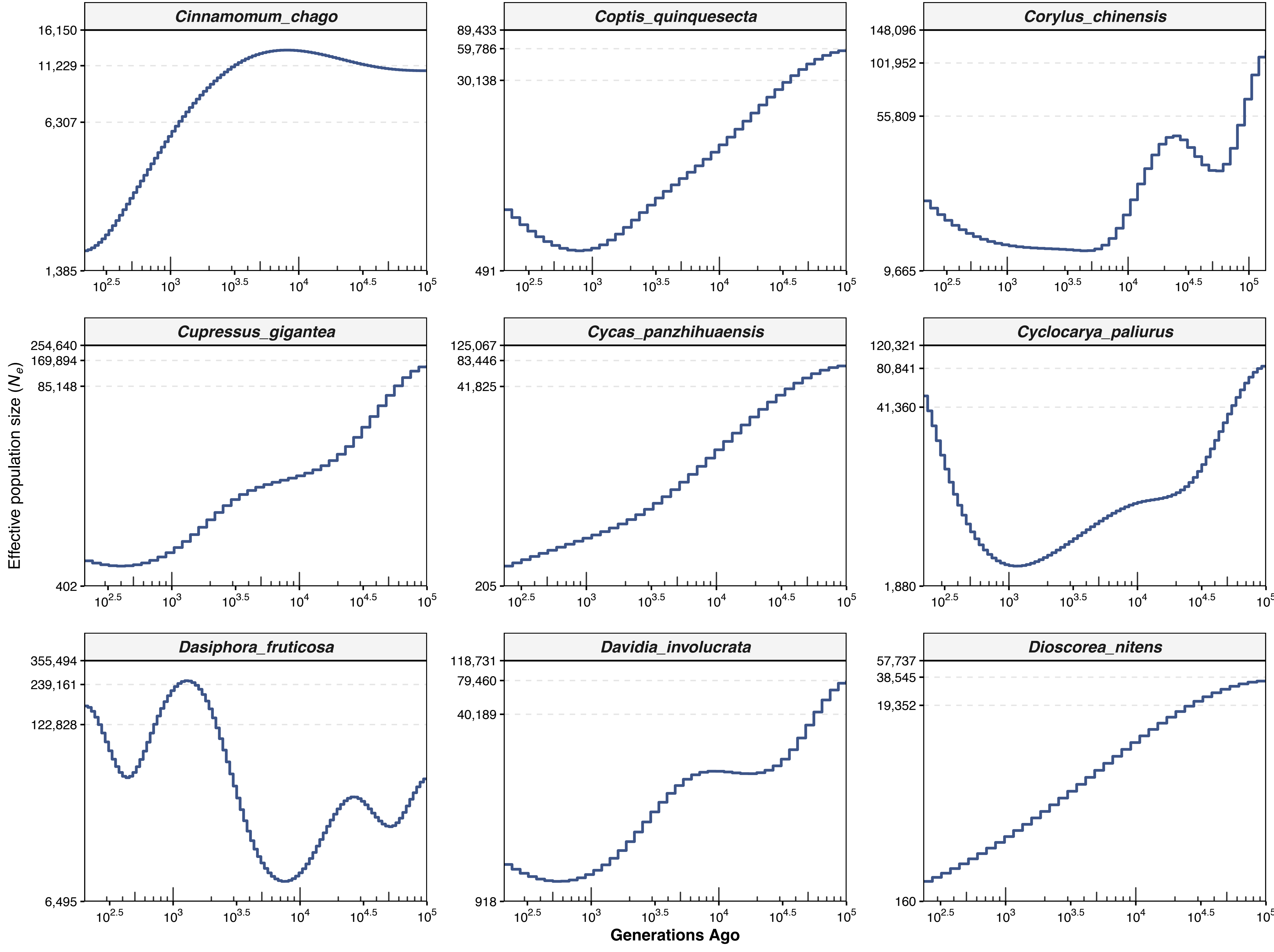

Supplementary Figure: Effective Population Size over Generations (Page 5 of 12 )

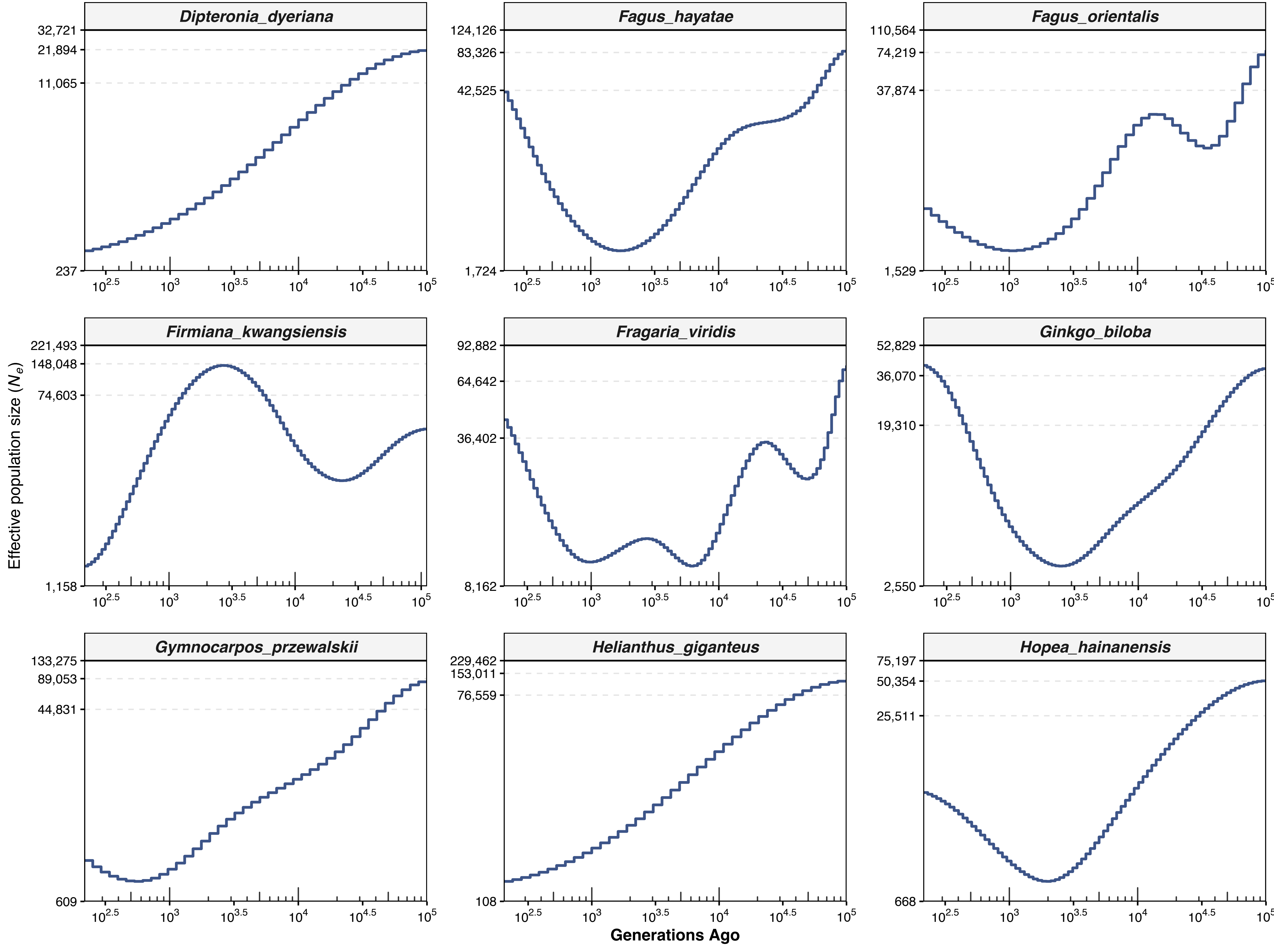

Supplementary Figure: Effective Population Size over Generations (Page 6 of 12 )

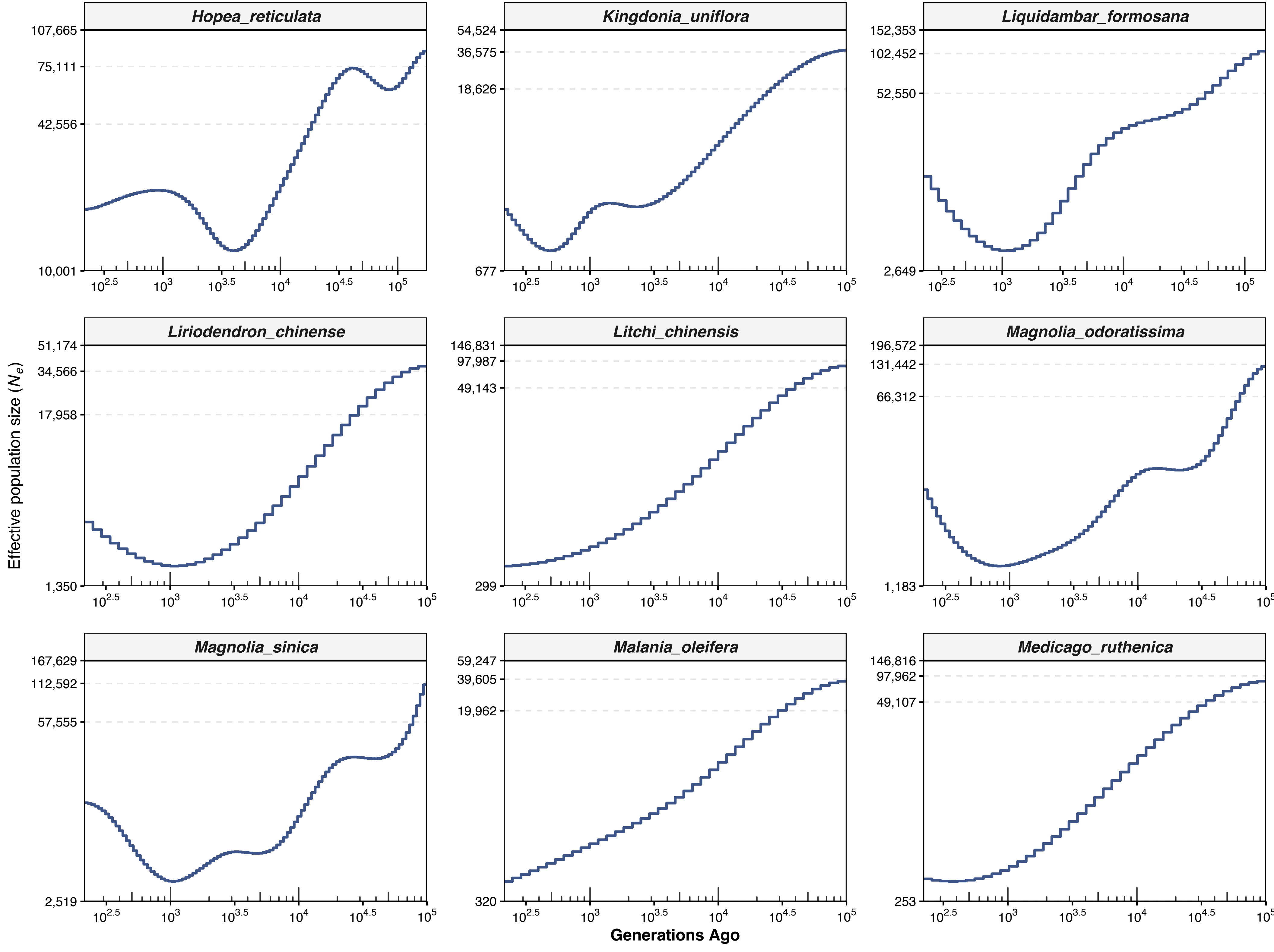

Supplementary Figure: Effective Population Size over Generations (Page 7 of 12 )

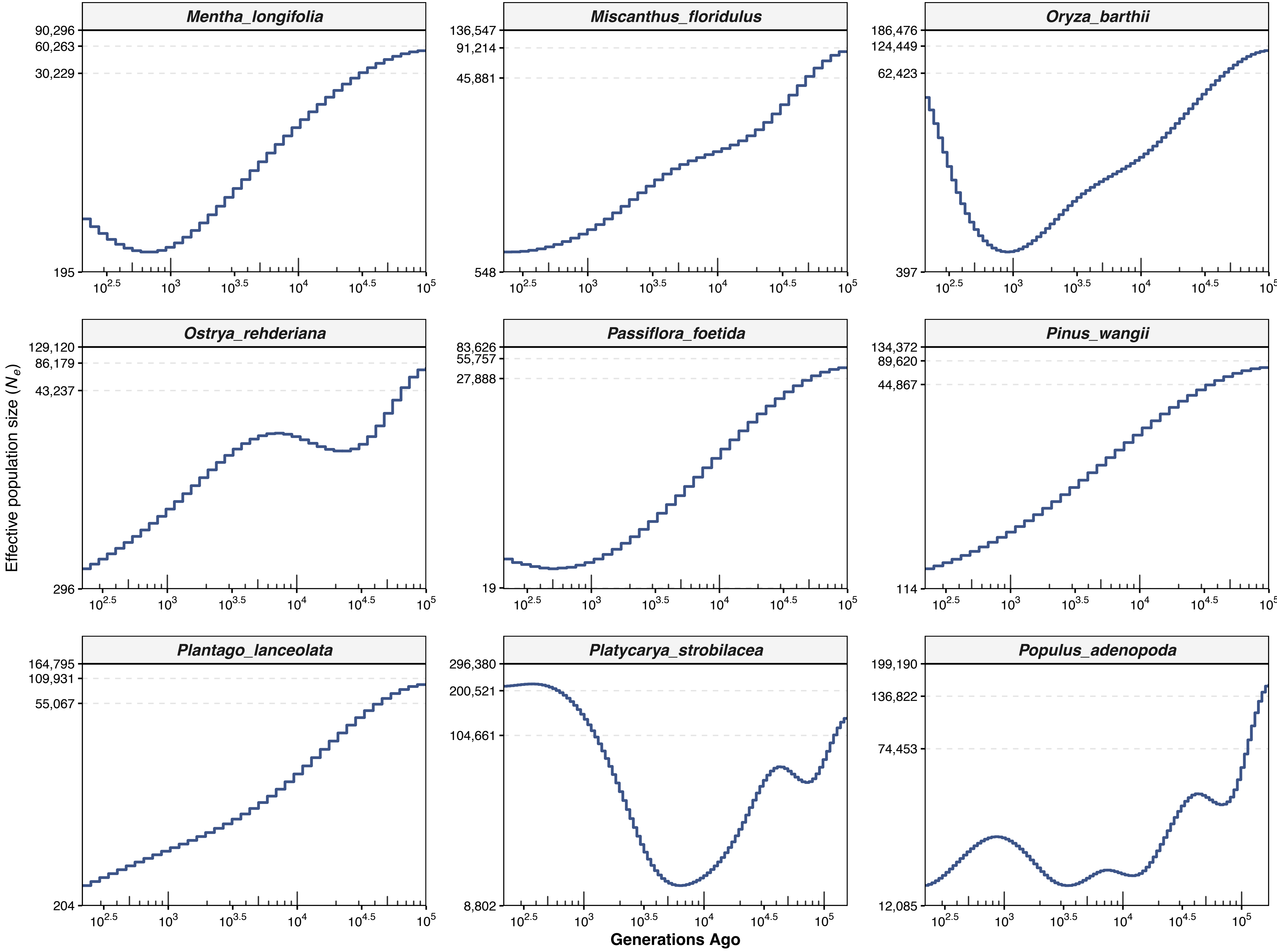

Supplementary Figure: Effective Population Size over Generations (Page 8 of 12 )

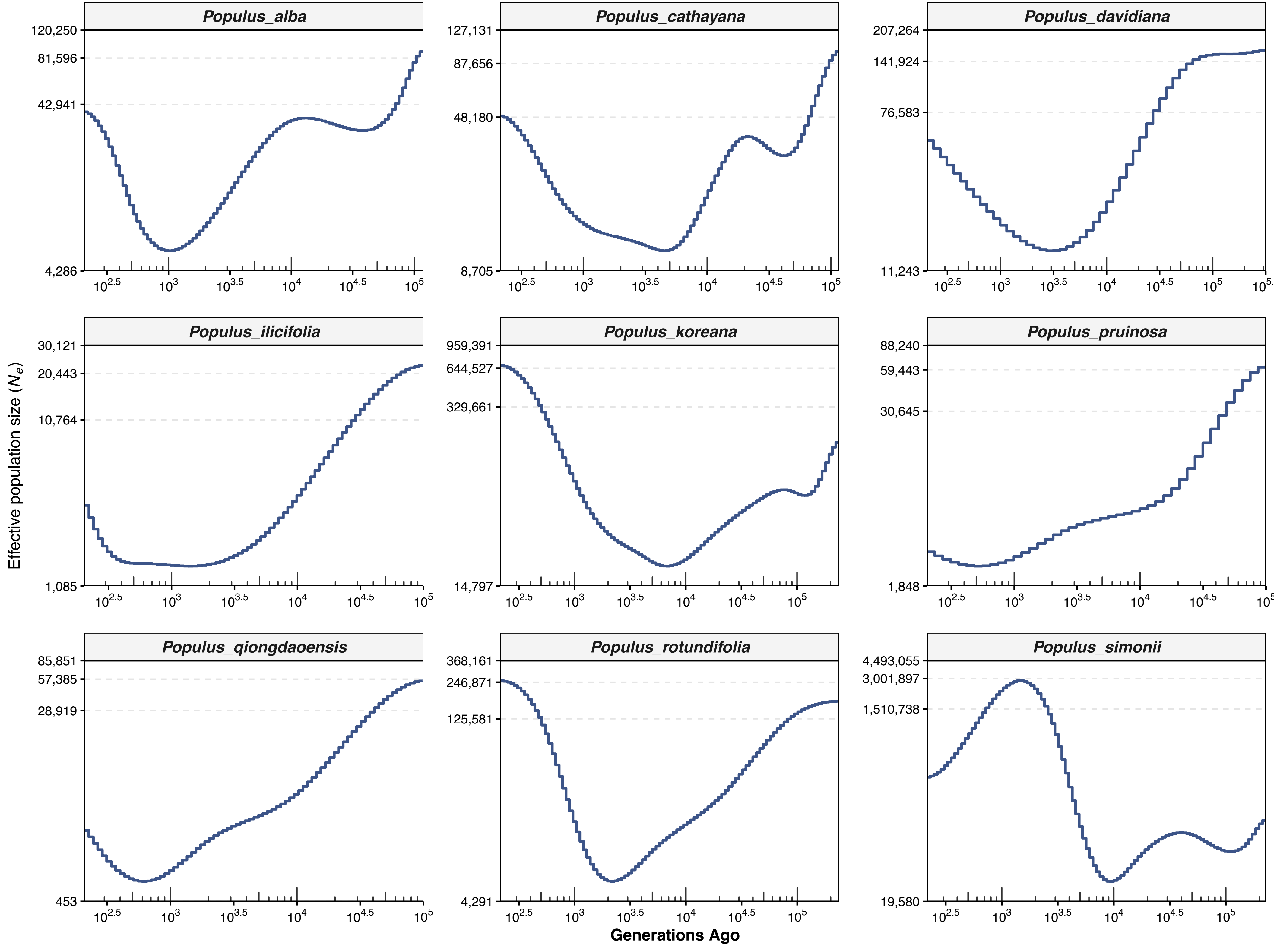

Supplementary Figure: Effective Population Size over Generations (Page 9 of 12 )

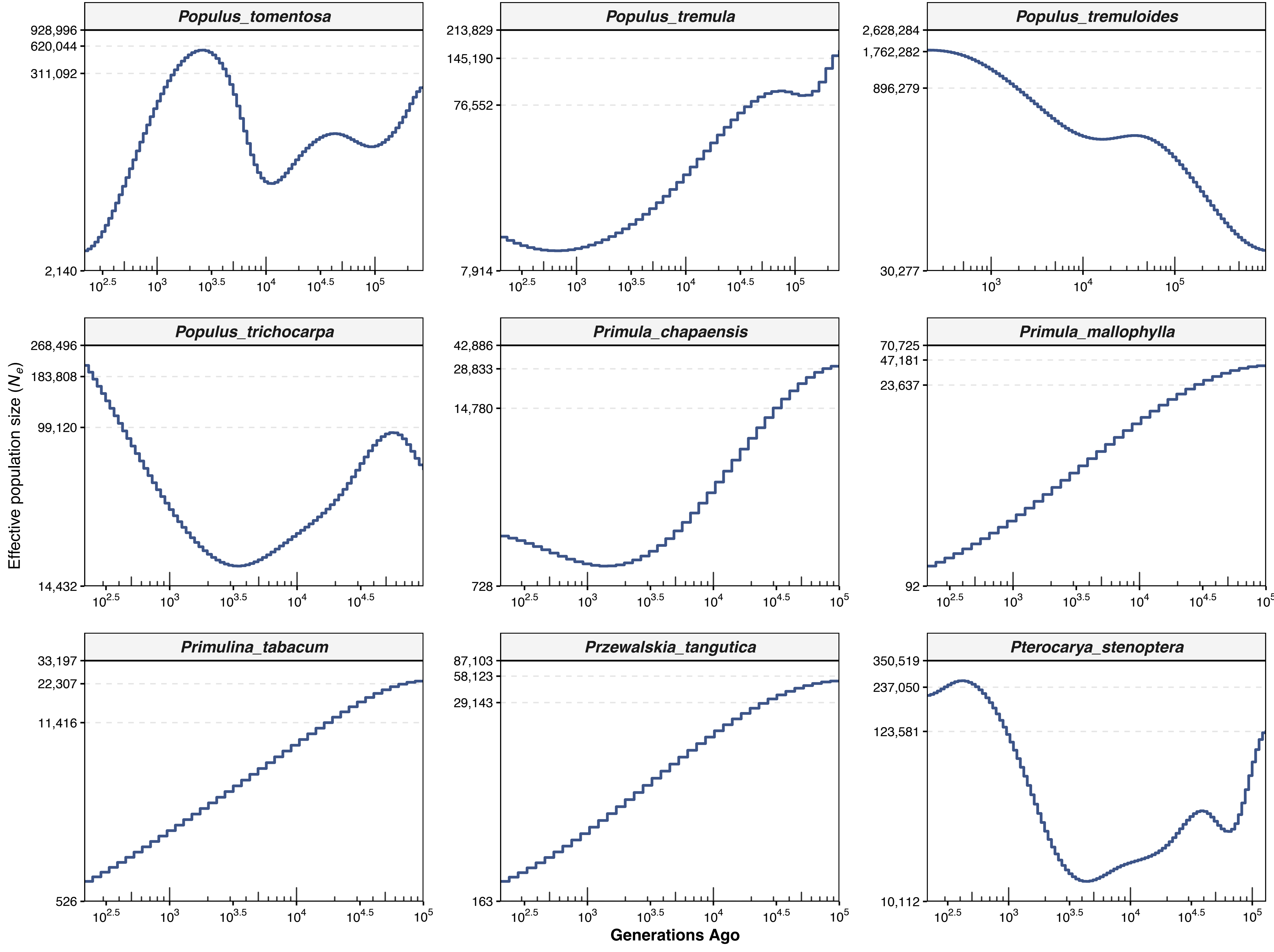

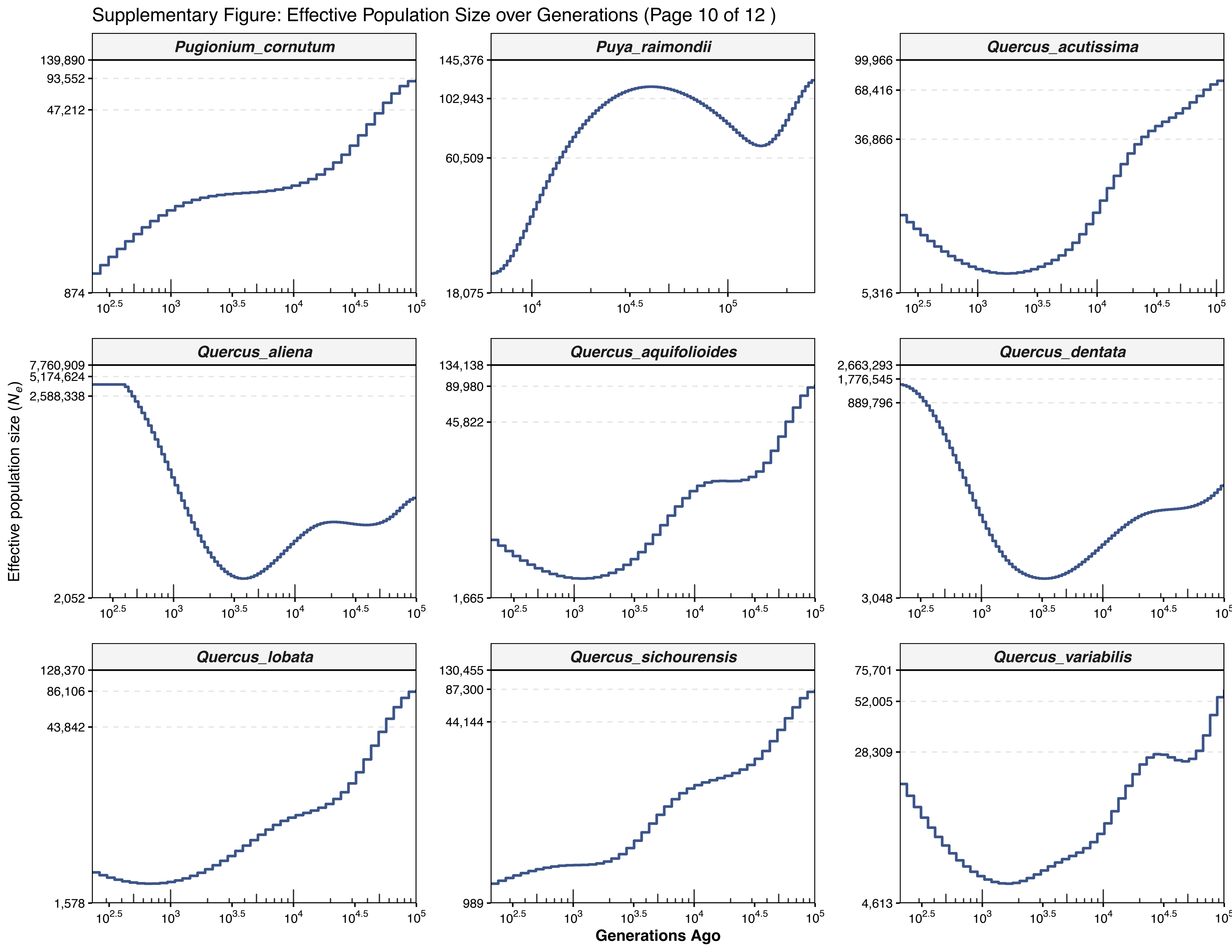

Supplementary Figure: Effective Population Size over Generations (Page 10 of 12 )
Pugionium_cornutum
Puya_raimondii
Quercus_acutissima
Quercus_aliena
Quercus_aquifolioides
Quercus_dentata
Quercus_lobata
Quercus_sichourensis
Quercus_variabilis
139,890
93,552
47,212
874
145,376
102,943
60,509
18,075
99,966
68,416
36,866
5,316
7,760,909
5,174,624
2,588,338
2,052
134,138
89,980
45,822
1,665
2,663,293
1,776,545
889,796
3,048
128,370
86,106
43,842
1,578
130,455
87,300
44,144
989
75,701
52,005
28,309
4,613
Effective population size ($N_e$)
Generations Ago

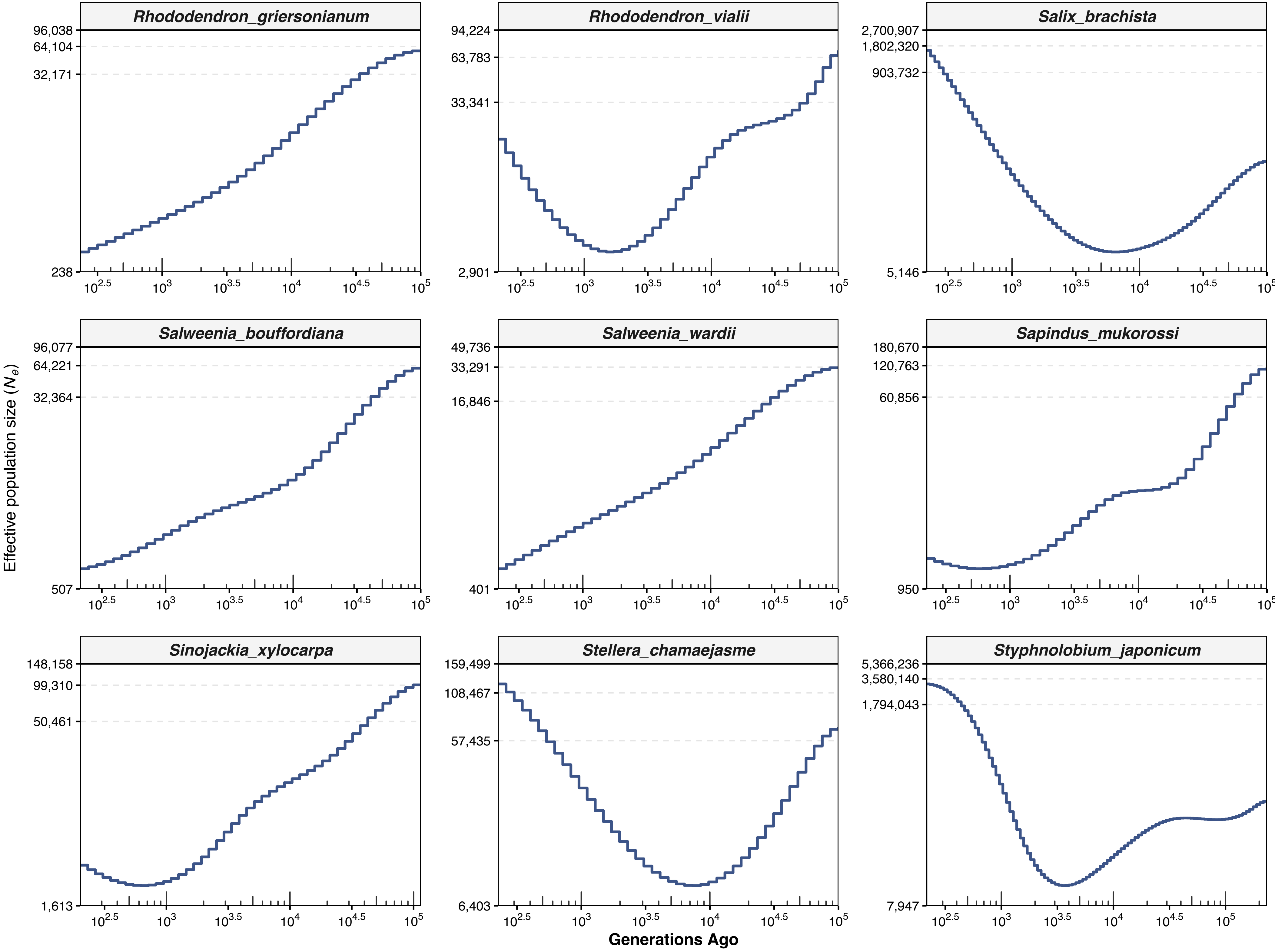

Rhododendron_griersonianum
96,038
64,104
32,171
238
Rhododendron_vialii
94,224
63,783
33,341
2,901
Salix_brachista
2,700,907
1,802,320
903,732
5,146
Salweenia_bouffordiana
96,077
64,221
32,364
507
Salweenia_wardii
49,736
33,291
16,846
401
Sapindus_mukorossi
180,670
120,763
60,856
950
Sinojackia_xylocarpa
148,158
99,310
50,461
1,613
Stellera_chamaejasme
159,499
108,467
57,435
6,403
Styphnolobium_japonicum
5,366,236
3,580,140
1,794,043
7,947
$10^{2.5}$ $10^3$ $10^{3.5}$ $10^4$ $10^{4.5}$ $10^5$
Effective population size ($N_e$)
Generations Ago

Supplementary Figure: Effective Population Size over Generations (Page 12 of 12 )

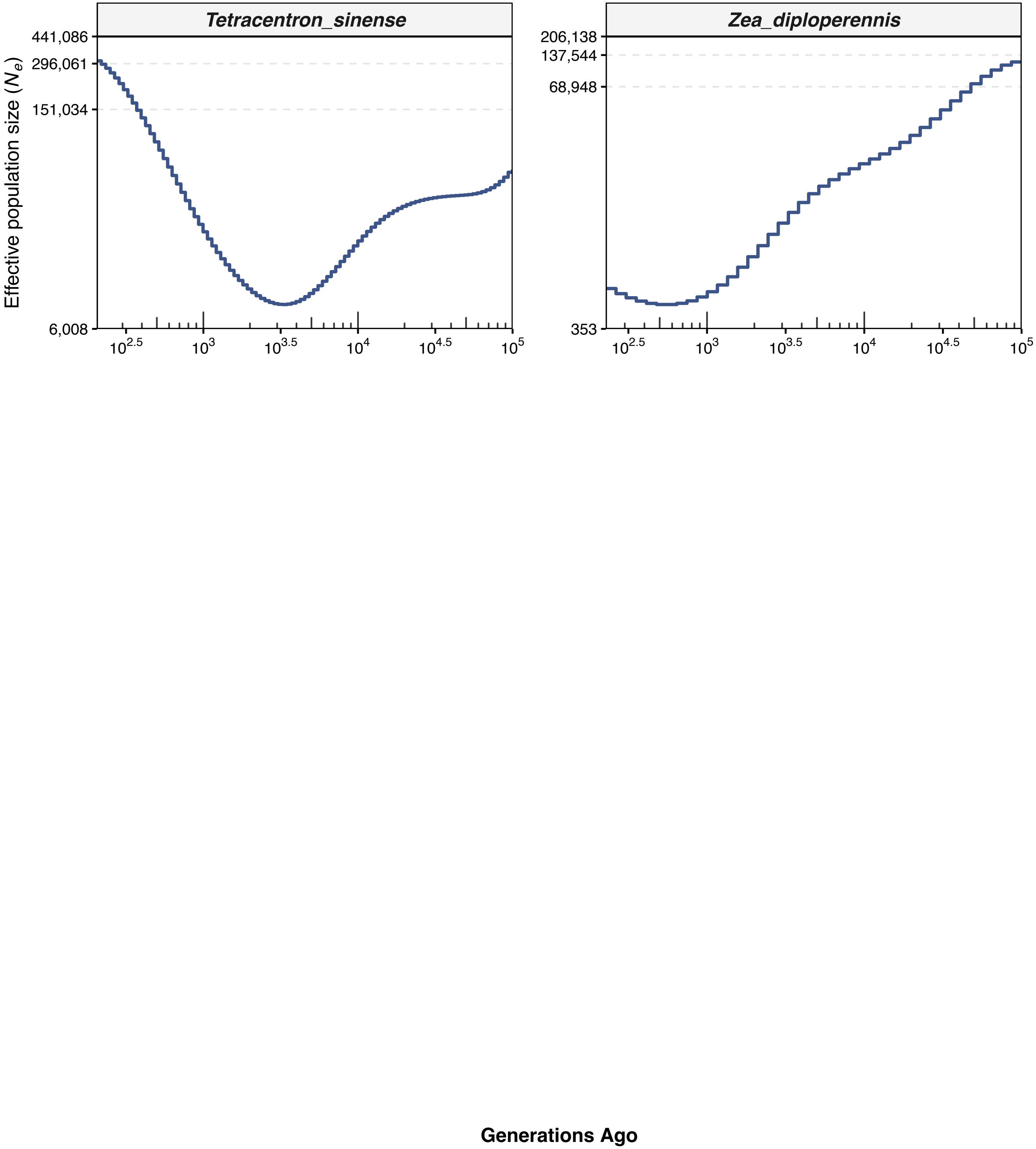
